# Dynamic switching of cell-substrate contact sites allows gliding diatoms to modulate the curvature of their paths

**DOI:** 10.1101/2025.03.18.643962

**Authors:** Stefan Golfier, Veikko F. Geyer, Nicole Poulsen, Stefan Diez

## Abstract

The directed motility of unicellular organisms is critical for their survival and ecological success, yet the mechanisms that enable rigid-walled cells to dynamically reorient and alter the shape of their trajectories remain poorly understood. Here, we investigate the gliding motility of *Craspedostauros australis*, a raphid pennate diatom that moves rapidly across submerged surfaces using an intracellular actomyosin motility complex and adhesive mucilage strand secretion. Using high-precision single-cell tracking, scanning electron microscopy (SEM), and interference reflection microscopy (IRM), we reveal how diatoms achieve diverse path curvatures by dynamically modulating the location of raphe-substrate contact and switching between one- and two-raphe contact gliding. Our results show that local curvature variations along the raphes dictate trajectory shapes, with one-raphe contact gliding producing highly curved paths, while two-raphe contact gliding results in paths of lower curvature. IRM imaging further confirms that transitions between these gliding modes underlie abrupt changes in path curvature and cell reorientation. This dynamic-raphe-switching mechanism is conserved across cell sizes and correctly predicts the increased path curvatures observed in smaller cells according to their more pronounced local raphe curvature. By quantitatively linking raphe geometry, cell-substrate attachment dynamics and motility patterns, our study provides insights into a novel motility mechanism that allows diatoms to adapt their movement to complex environments.

## Introduction

Motility is a fundamental hallmark of life across all scales. From single cells to large organisms, diverse mechanisms have evolved to traverse space in search of food, mating partners and favorable conditions while evading predation and hazardous environments. Free-swimming single-celled organisms often rely on beating cilia and flagella (as in some green algae or sperm cells) or rotating flagella (as in some bacteria) that allow for speeds of several 100 µm/s (Milo & Phillips, 2015). In contrast, surface-bound motility such as amoeboid crawling, apicomplexan gliding as well as bacterial twitching and gliding, is slower, with speeds typically below 1 µm/s (McBride, 2001). An intriguing exception are diatoms, an abundant class of unicellular microalgae encased in a rigid silica cell wall, many of which exhibit rapid gliding motility of up to 35 µm/s along complex trajectories on submerged surfaces (Edgar, 1979). Remarkably, they achieve this without extracellular appendages or cell deformation (Edgar & Pickett-Heaps, 1984; Bertrand, 1990).

Diatoms are broadly classified into two major groups based on cell shape: radially symmetric ‘centric’ diatoms, which exist mainly as free-floating phytoplankton, and bilaterally symmetric ‘pennate’ diatoms, which are predominantly benthic, colonizing sunlit underwater surfaces. A subgroup of pennate diatom species is highly motile, which is enabled by a system of longitudinal slits in their cell wall, termed ‘raphes’. These raphid pennate diatoms adhere to surfaces via the secretion of adhesive mucilage strands through the raphe (Drum & Hopkins, 1966; Edgar, 1983; Harper & Harper, 1967; Higgins et al., 2003; Wang et al., 2000). The forces for motility are likely produced by an intracellular actomyosin force-generating motor translocating the substrate-bound adhesive mucilage strands along the raphe in order to propel the cell forward, termed the adhesion motility complex (AMC) (**Figure 1A**, for review see (Edgar & Pickett-Heaps, 1984; Poulsen et al., 2022). Despite numerous theories, the exact mechanism of diatom propulsion remains controversial.

**Figure 1:**
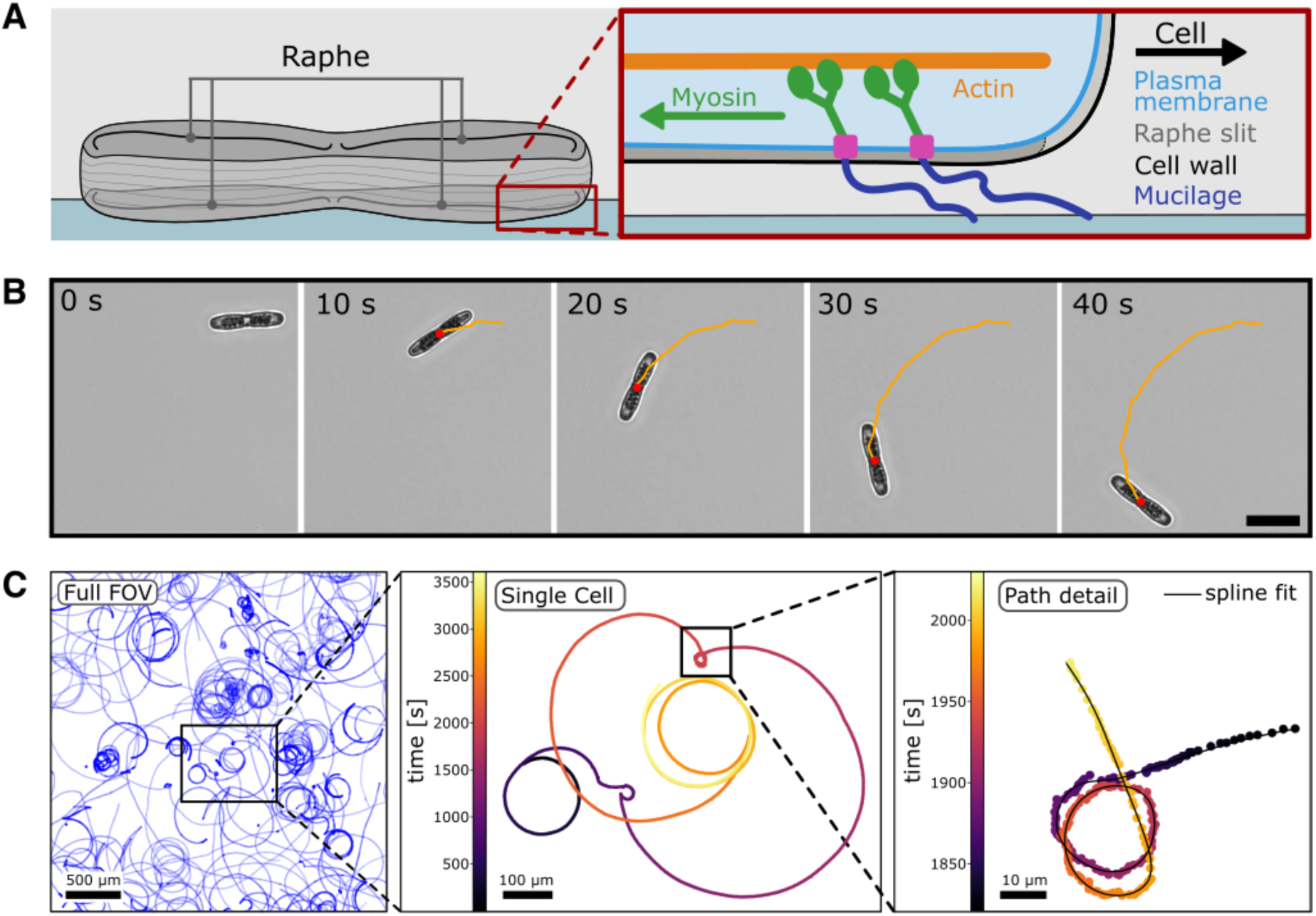
Single-cell tracking of motile *C. australis*. **A** Schematic representation of a diatom gliding on a glass surface. Inset: Current understanding of the diatom adhesion motility complex (AMC). Intracellular myosin motors (green) couple via a transmembrane complex (pink) to adhesive mucilage strands (dark blue), which are secreted through the raphe in the cell wall (grey) and adhere to the glass surface. Cell motility is achieved through myosin movement on intracellular actin cables (orange), leading to tension on the adhesive mucilage strands that propels the cell forward. **B** Selected timelapse images (acquired at 3 fps) of the motility of an individual diatom cell over 40 s. The trajectory (orange) is derived from high-precision tracking of the center of mass (red) in each image. **C** Trajectories of multiple *C. australis* cells over one hour (acquired at 1 fps). Left panel: Full field of view, displaying over 100 individual single-cell trajectories. Mid panel: Typical trajectory of an individual cell displaying several changes in path curvature. Color-code describes time. Right panel: Zoom-in into the same track illustrating the spatial and temporal resolution employed in this study as well as a spline fit used to quantify path curvature.

The ecological success of raphid pennate diatoms in their complex and dynamic environments can in large parts be attributed to their active motility, enabling them to colonize new aquatic habitats, optimize light exposure (S. A. Cohn et al., 1999, 2015, 2016; Duchêne et al., 2023; Jesus et al., 2023; McLachlan et al., 2009; Morelle et al., 2024), enhance nutrient uptake (Bondoc, Heuschele, et al., 2016; Cooksey & Cooksey, 1988; V. Bondoc et al., 2019) and pursue reproductive strategies (Bondoc, Lembke, et al., 2016). The pronounced photo- and chemotactic motility of diatoms (Apoya-Horton et al., 2006; Nultsch, 1971) is based on their ability to navigate along strikingly complex paths, featuring dynamic changes in path curvature and directional reversals, allowing them to quickly reorient their movement (Bondoc et al., 2016). However, how cells that are encased in a rigid cell wall achieve dynamic directional flexibility without cell deformation and extracellular appendages remains poorly understood.

Several studies have proposed that the shape of a diatom’s trajectory is governed by the geometry of its raphe system (Nultsch, 1956; Round, 1990; S. A. Cohn & Weitzell, 1996; S. A. Cohn, 2001; Harbich, 2021; Bondoc-Naumovitz et al., 2024). Specifically, it has been suggested that traction is exerted over only a few micrometers of the raphe system in contact with the substrate at any given time and that the local curvature of this active zone dictates the cell’s trajectory at that moment (Harbich, 2021). By shifting this active zone along the raphe and thereby engaging a segment of different curvature, diatoms may be able to dynamically adjust their path curvature, enabling directional flexibility despite their rigid cell wall. However, a mechanistic understanding of how local raphe geometry quantitatively determines trajectory shape and how transitions between different path curvatures occur remains lacking.

Here, we propose and experimentally validate a mechanism, that quantitatively links raphe geometry, cell-substrate attachment dynamics and cell size to the trajectories of *Craspedostauros australis*, a model species for studying diatom adhesion and motility (Lind et al., 1997; Abbriano, 2023). Using high-precision single-cell tracking, scanning electron microscopy (SEM) and interference reflection microscopy (IRM), we find that the observed spectrum of path curvatures can only be explained by distinct modes of gliding motility, each enabling traction in different regions of the raphe. Direct correlations between cell-substrate contact sites and path curvature in IRM confirm, that cells dynamically transition between these modes, leading to abrupt changes in path curvature and cell reorientation. Based on these findings, we propose a new model of diatom motility that quantitatively predicts the observed cell-size dependency of *C. australis* trajectories and advances our understanding of cellular movement.

## Results

### Single-cell tracking of motile diatoms reveals dynamic changes in path curvature

To investigate the motility characteristics of *C. australis*, we settled cells grown in artificial seawater medium onto hydrophobically-coated glass coverslips and recorded time-lapse movies using brightfield microscopy (**Figures 1B, Methods**). Each field of view (3.2 x 3.2mm) contained up to 100 cells, which were imaged for time periods of up to one hour with a frame rate of 1 frame-per-second (fps) (**Supplementary Movie S1)**. For every frame, we extracted the spatial coordinates of each cell by determining the position of its center of mass, allowing us to reconstruct trajectories of individual cells (**Figure 1C**). Over short timescales (tens of seconds to minutes), the trajectories of single cells typically followed circular paths with conserved radii, as was previously observed in other raphid diatoms (Edgar & Pickett-Heaps, 1984; Gutiérrez-Medina et al., 2014). However, over longer timescales, individual cell trajectories exhibited a remarkable variability in their path curvatures **(Supplementary Movie S2)**. We frequently observed abrupt and substantial changes in path curvature, both from quasi-straight to highly curved paths and vice versa. Additionally, we noticed a strong variability in the overall shapes of the trajectories between different cells (**Supplementary Figure S1**). This is surprising, because we studied cells derived from mono-clonal cell cultures (i.e. with similar raphe morphology) and performed experiments under isotropic conditions (i.e. under homogeneous illumination, homogeneous treatment of the substrate surface and in the absence of chemical gradients). In light of previous literature suggesting raphe geometry as the basis of path shape (Nultsch, 1956; Round, 1990; S. A. Cohn & Weitzell, 1996; S. A. Cohn, 2001; Harbich, 2021; Bondoc-Naumovitz et al., 2024), we were wondering about the source of heterogeneity in path shapes from a cell population that should feature similar raphe geometries. We thus sought to identify potential correlations between the motility parameters that would allow us to formulate testable hypotheses on the underlying mechanisms.

### Path curvature is correlated with cell velocity and orientation

To identify potential correlations between the motility parameters, we quantitatively analyzed the path curvature and the velocity of each individual cell along its trajectory, as well as the cell’s orientation with respect to its direction of travel (see **Figure 2 A-C** and Methods for the respective definitions). For simplicity, we first discuss our results for an exemplary trajectory of a single cell tracked over 890 s (approximately 15 min), during which the cell traversed a total distance of 2400 µm (**Supplementary Movie S3)**. Initially the cell followed a trajectory with low curvature (curvature of about 0.001 µm^-1^). After 450 µm along the path, the path curvature suddenly increased by more than two orders of magnitude (**Figure 2A**, right panel vertical dashed grey line). Over the next 2000 µm, the path curvature gradually decreased, resulting in a spiral-shaped trajectory with increasing radius. The velocity profile of this trajectory reveals that the cell significantly slowed down before transitioning from a path with low curvature (quasi-straight) to a path with high curvature (tight circles) (**Figure 2B**). Afterwards, it gradually accelerated as the path curvature decreased, regaining its initial velocity once the path curvature reached the initial level. Furthermore, we found that along path segments of high curvature, the orientation of the cell’s long axis displayed a significant offset (up to 60 degrees) with respect to the direction of travel (**Figure 2C, Supplementary Movie S4**). Closer inspection revealed that just before the sudden increase in path curvature, the back of the cell started to spin around its leading apex, resulting in an abrupt increase in the cell’s offset angle. This behavior we observed for different cells upon stark increase in path curvature (**Supplementary Movie S5**).

**Figure 2:**
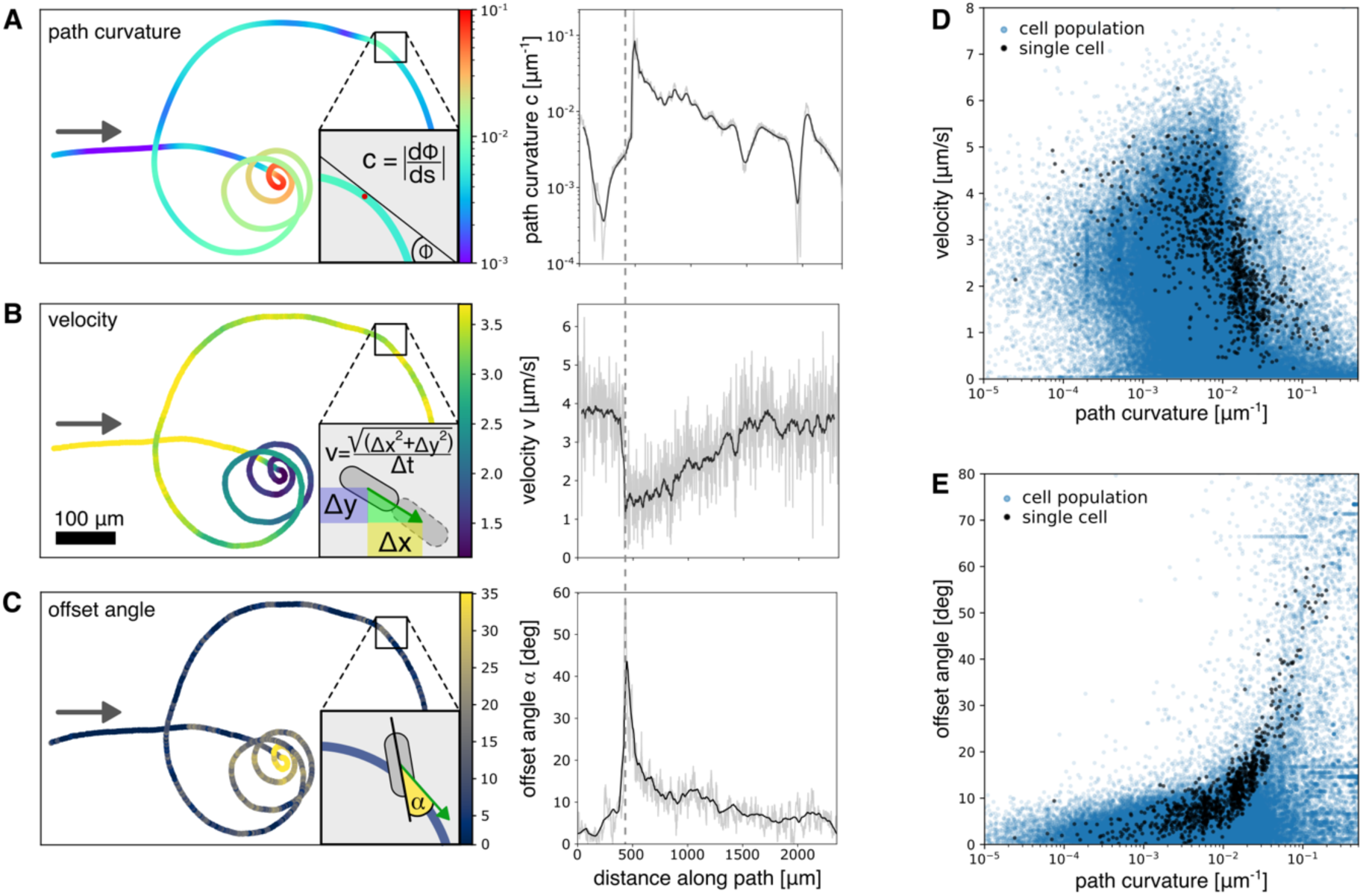
Parameters to quantify the motility of *C. australis* cells. **A** Quantification of path curvature c for a single cell trajectory. Gray arrow indicates the direction of travel. Inset depicts the definition of path curvature (Methods). Color-code describes path curvature. (right) Path curvature along the path with raw data in light gray, rolling average with window size of 4 µm in black. **B** Quantification of cell velocity v (same track as in A). Inset depicts the definition of velocity along the path with the velocity vector in green (Methods). Color-code describes cell velocity. (right) Cell velocity along the path with raw data in light gray and rolling average with window size of 4 µm in black. **C** Quantification of offset angle α (same track as in A). Inset depicts the definition of the offset angle (yellow) as the absolute difference between the orientation of the cells’ long axis (black line) and the direction of travel (green arrow) (Methods). Color-code describes offset angle. (right) Offset angle along the path with raw data in light gray and rolling average with window size of 4 µm in black. **D** Correlation between cell velocity and path curvature for the single cell track from A-C (black dots) and a cell population of 196 cells (blue dots) over timescales of up to one hour. **E** Correlation between offset angle and path curvature for the same single cell track (black dots) and the same cell population (blue dots).

When quantitatively relating the measured motility parameters to each other, we found that cell velocity was anti-correlated to path curvature (Spearman’s rank correlation coefficient of −0.73), and decreased exponentially with increasing path curvature (**Figure 2D** grey dots). Consequently, the cell displayed maximum velocities (up to 8 µm/s) along quasi-straight path segments (curvature < 0.001 µm^-1^) and slowed down significantly once the curvature increased above 0.03 µm^-1^. In contrast, the cells’ offset angle was correlated with path curvature (Spearman’s rank correlation coefficient of 0.85), with large offset angles along highly curved path segments and small offset angles along quasi-straight path segments (**Figure 2E** grey dots). Conversely, cell velocity decreased exponentially with increasing offset angle (**Supplementary Figure S2A**). These relationships between path curvature, cell velocity, and offset angle, are, albeit more heterogeneous, also observed on a population level (**Figure 2D and E** blue dots, 196 cells tracked over timescales of up to one hour, showing a characteristic velocity peak around 2 µm/s, **Supplementary Figure S2B**). These data indicate, that it is unlikely for a cell to display rapid movement along stretches of high path curvature and similarly unlikely for a cell to show high offset angles while moving along path sections of low curvature.

Taken together, our observations demonstrate, that the path curvature of motile *C. australis* cells is highly variable and strongly correlates with cell velocity and offset angle. In particular, the high offset angles in path segments with high curvature suggest that traction is generated along short, discrete segments of the raphe, leading to low forces for motility (and hence lower velocities) with the local raphe curvature determining the path curvature. On the other hand, the low offset angles observed along segments with low path curvature suggest, that longer sections of the raphe engage with the substrate, enabling greater forces for motility (and hence higher velocities). As such, our findings support the hypothesis that the motility of diatom cells might be governed by short segments of the raphe system in contact with the substrate (Harbich, 2021). However, the highly dynamic nature of the observed trajectories contrasts with the structural rigidity of the raphe system, underscoring the need for a dynamic mechanism that translates static local raphe geometry into adaptable path curvatures.

### Path curvature can be predicted from raphe curvature by distinct modes of gliding motility

To test the hypothesis that raphe curvature governs path curvature, we sought a way to quantitatively compare the spectrum of local raphe curvatures of a cell population to the spectrum of path curvatures from cells of the same population. To that end we preserved cells of the population used to generate Figure 2D and 2E and performed SEM imaging of their silica cell walls. **Figure 3A** depicts an SEM image of a typical *C. australis* cell wall with the two raphes visible as dark ridges running along the length of the cell, interrupted by a silica bridge in the middle of the cell (central nodule). We manually traced each pair of raphes per cell in the SEM images and used the obtained *x-y*-data to quantify the local raphe curvature (**Figure 3B, Supplementary Figure S3A, Methods**). Specifically, we segmented the raphes of 20 cells and analyzed the local raphe curvature within a 4 µm window, which we shifted along the length of the raphe. We found, that the local curvature along the raphes of each cell varies over several orders of magnitude, with highly curved regions at the cells’ ends (terminal raphe fissures) and centers (close to the central nodule) as well as stretches of low curvature in between (**Figure 3B)**. In all cases the shapes of the raphes followed the bilateral symmetry of the cell, displaying similar curvature patterns in both halves of the cell.

**Figure 3:**
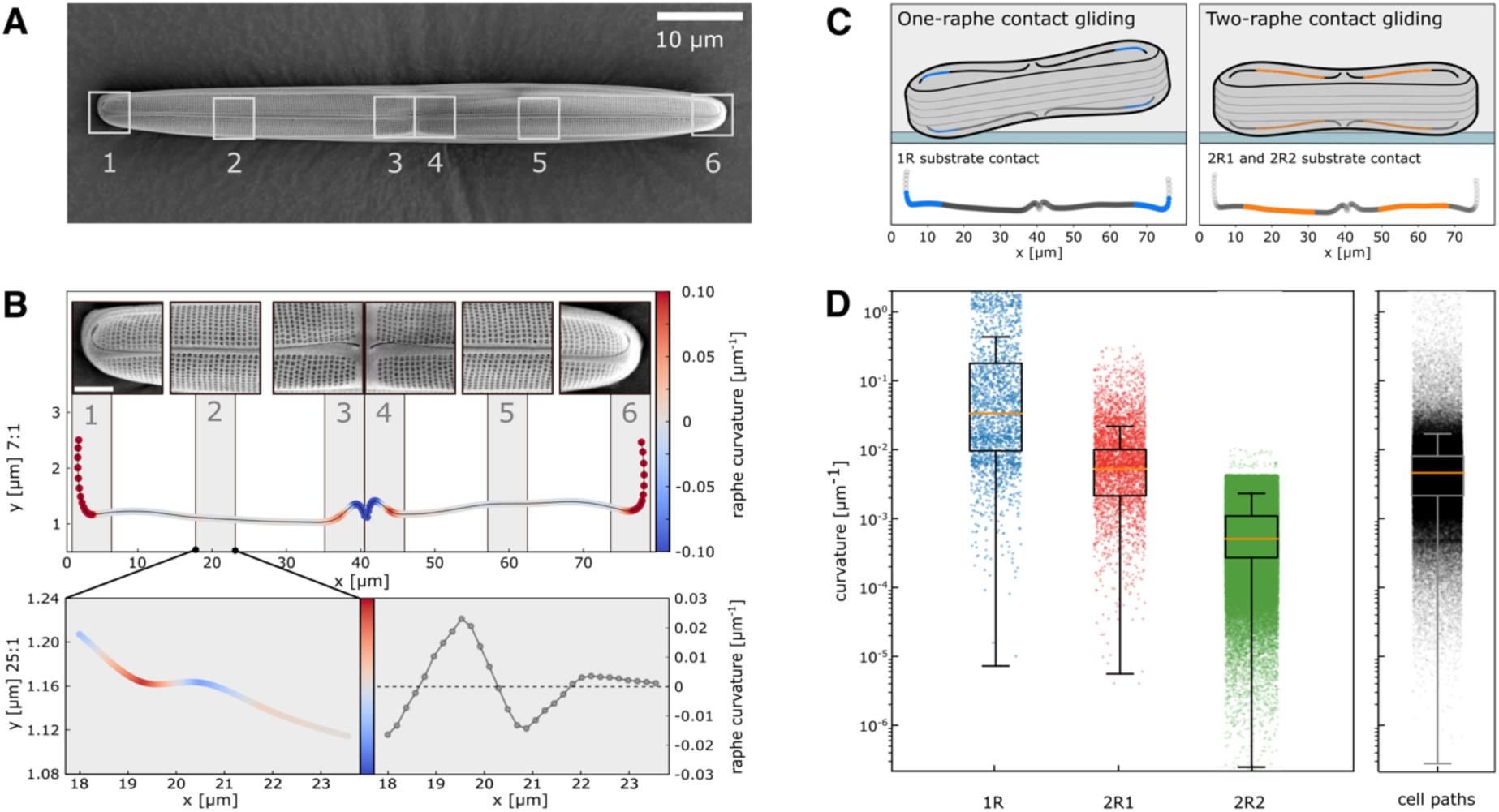
Quantification of *C. australis* local raphe curvature. **A** Scanning electron microscopy (SEM) image of the silica cell wall of one *C. australis* cell. **B** Quantification of raphe curvature along the two raphes in A, with regions of interest (1-6) to illustrate the shape of the raphe. (bottom) Zoom-in into a central section of the raphe to illustrate the local variability of the raphe curvature. **C** Modes of diatom gliding motility. (left) One-raphe contact gliding (1R) requires the cell to pivot up (away from the substrate) in order to allow only a single stretch of one raphe to contact the substrate. Cell wall geometry restricts the regions of contact with the substrate to about the terminal third of each raphe, illustrated in blue. The terminal 1 µm in each raphe was excluded as it is filled with silica. (right) During two-raphe contact gliding the cell makes simultaneous contact to the substrate with two segments along the raphes in both halves of the cell. Cell wall geometry restricts the regions of substrate contact to the central half of each raphe, highlighted in orange. Forces for gliding motility may be produced along only one segment in one of the two raphes (2R1) or along two segments in both raphes simultaneously (2R2). The central nodule and its vicinity can never make contact to the substrate as the cell wall curves inwards in these regions and was thus excluded from analysis. **D** (left) Spectrum of local raphe curvatures along 4 µm windows in the one-raphe (blue) and two-raphe (red) contact zones, representing the predicted path curvatures for 1R and 2R1 gliding respectively. Predicted path curvatures for two-raphe contact gliding with simultaneous activity on both raphes (2R2) were obtained from circle-fits to all possible combinations of the two-raphe contact zones in both raphes (green). Local raphe curvatures were quantified using SEM images of 18 diatom cell walls, prepared from the cell population in Figure 2. (right) Spectrum of experimentally observed path curvatures from 196 trajectories of the cell population in Figure 2, averaged over 4 µm windows along the path.

If path curvatures of a motile diatom were to be predicted from the local geometry of its raphe, one needs to consider which parts of the raphe could actually contribute to gliding and thus govern path curvature. For this, we assumed that only parts of the raphe that can make direct contact to the substrate can create traction for motility, as has been suggested by recent observations (Harbich, 2021; Davutoglu et al., 2024; Bondoc-Naumovitz et al., 2024). Our SEM images of *C. australis* cell walls revealed that the geometry of the cell wall restricts which regions are in direct contact with the substrate (**Supplementary Figure S3B**). Based on this, we subdivided the raphes into contact zones. We found that the terminal 30% of each raphe can only touch the substrate, if the cell pivots up (away from the substrate). This leads to only one zone of the raphe to come into contact with the substrate - a scenario which we denote as “one-raphe contact gliding” (1R) (**Figure 3C** left panel, **Supplementary Figure S3B and C** blue region and dots). Quantitative analysis of the local raphe curvatures in these regions revealed a median curvature of about 0.04 µm^-1^ (**Figure 3D** left panel 1R). As in the case of one-raphe contact gliding, only a single segment of one raphe can generate traction at any time, we assume that the measured raphe curvature directly provides a prediction of the expected path curvature.

In contrast, when the cell lies flat, the central 50% of each raphe simultaneously make contact with the substrate - a scenario we denote as “two-raphe contact gliding” (2R) (**Figure 3C** right panel, **Supplementary Figure S3B and C** orange region and dots). However, it remains unclear whether traction is generated along only one raphe (2R1) or in both raphes (2R2) simultaneously. If force generation occurs only along a single zone of one raphe, we again assume that the measured median raphe curvature of about 0.007 µm^-1^ (pooled curvature values in the two-raphe contact zones) directly provides a prediction of the expected path curvature (**Figure 3D** left panel 2R1). This value is already significantly lower than for one-raphe contact gliding, highlighting the fact, that the raphe segments which can produce traction during two-raphe contact gliding are less curved than the raphe segments that exclusively drive one-raphe contact gliding.

However, recent studies using the same diatom species showed concerted activity of myosin motors in both raphes during smooth gliding (Davutoglu et al., 2024). This suggests a mode of diatom gliding in which forces for motility are produced in both raphes simultaneously. Consequently, we extended the idea of an individual raphe zone determining the path curvature suggested in literature (Harbich, 2021; Zhang et al., 2024; Bondoc-Naumovitz et al., 2024) and hypothesized that during two-raphe contact gliding two contact zones (one in each raphe) can also simultaneously contribute to cell motility and hence influence path curvature. The ensuing path curvature will then be a combination of the local raphe curvatures of the zones in contact with the substrate and their relative distance to each other (see **Methods** for precise description of the analysis). Predicting path curvatures in this way yielded significantly lower values, suggesting nearly straight trajectories during this mode of gliding (median predicted path curvature of about 0.001 µm^-1^, **Figure 3D** left panel 2R2).

Finally, by analyzing the path curvatures within 4 µm windows along the 196 diatom trajectories used to generate Figure 2D and 2E, we obtained a distribution of path curvatures with a median of 0.005 µm^-1^ (**Figure 3D** right panel). Comparing this distribution of cellular path curvatures to the distribution of path curvatures predicted from the local raphe geometry, we found that the predicted path curvatures during one-raphe contact gliding (1R) mapped only to high-curvature path segments, and failed to account for the bulk of the path curvatures below 0.01 µm^-1^. Conversely, the predicted path curvatures during two-raphe contact gliding (2R1 and 2R2) corresponded to the lower half of the path curvature spectrum, but failed to explain the highly curved path segments frequently observed in the cells’ trajectories. Specifically, simultaneous activity in both raphes (2R2) leads to the lowest predicted path curvatures, which can account for the lower end of the observed path curvatures where the cells moved along nearly straight segments. Taken together our data indicate, that only the combination of one-raphe and two-raphe contact gliding can account for the full range of path curvatures observed in our cell tracking data, suggesting that dynamic changes in path curvature originate from cells switching between these modes of gliding. To directly test whether changes in path curvature result from cells switching from one-raphe to two-raphe contact gliding, we next sought a method to experimentally quantify cell-substrate attachment dynamics in concert with path curvature.

### Substantial changes in path curvature result from switching between one- and two-raphe contact gliding

To examine if we can directly observe instances where gliding cells switch from one-raphe to two-raphe contact gliding and how this would affect path curvature, we employed interference reflection microscopy (IRM) to detect regions of cell-substrate contact while tracking the motility of the cells. In brief, IRM uses interference effects between light reflected from the coverslip and the cell wall. The intensity of the obtained signal is sensitive to the distance between the cell and the coverslip surface and can thus be used as a proxy for cell-substrate contact. This method allowed us, to directly track the position of the cell, analyze the curvature of its trajectory and correlate this with the number of raphes segments in contact with the substrate. **Figure 4A** depicts a trajectory from a single cell, with average path curvature values fluctuating over three orders of magnitude (**Figure 4C, Supplementary Movie S6**). Analyzing this and similar trajectories (**Supplementary Figure S4, Supplementary Movie S7)**, we found, that *C. australis* cells occasionally lost and re-established contact with the substrate with one raphe and hence switched between one-raphe and two-raphe contact gliding. A representative example is shown in **Figure 4B**, where in panel 1, only one end of the cell is in contact with the surface (black and white striated signal in red circle, indicating 1R contact) and in panel 2 both ends of the cell are in contact with the surface (black and white striated signal in red circles, indicating 2R contact). The resulting trajectories showed substantial increases in path curvature, when contact was lost in one raphe, and substantial reductions in path curvature once substrate contact with both raphes was re-established (**Figure 4C**). Analysis of all 11 trajectories that displayed switching between one- and two-raphe contact gliding revealed, that the path curvature during one-raphe contact gliding was an order of magnitude higher (median of 0.033 µm^-1^) than during two-raphe contact gliding (median of 0.002 µm^-1^) (**Figure 4D**). From this we conclude, that the substantial changes in path curvature originate from switches between one-raphe and two-raphe contact gliding. Interestingly, we observed that during switches from two-raphe to one-raphe contact gliding, traction can be lost in either the leading or the trailing raphe (see **Supplementary Movie S6, S7** and **S8)**. This suggests that the mechanism may be independent of the direction of travel.

**Figure 4:**
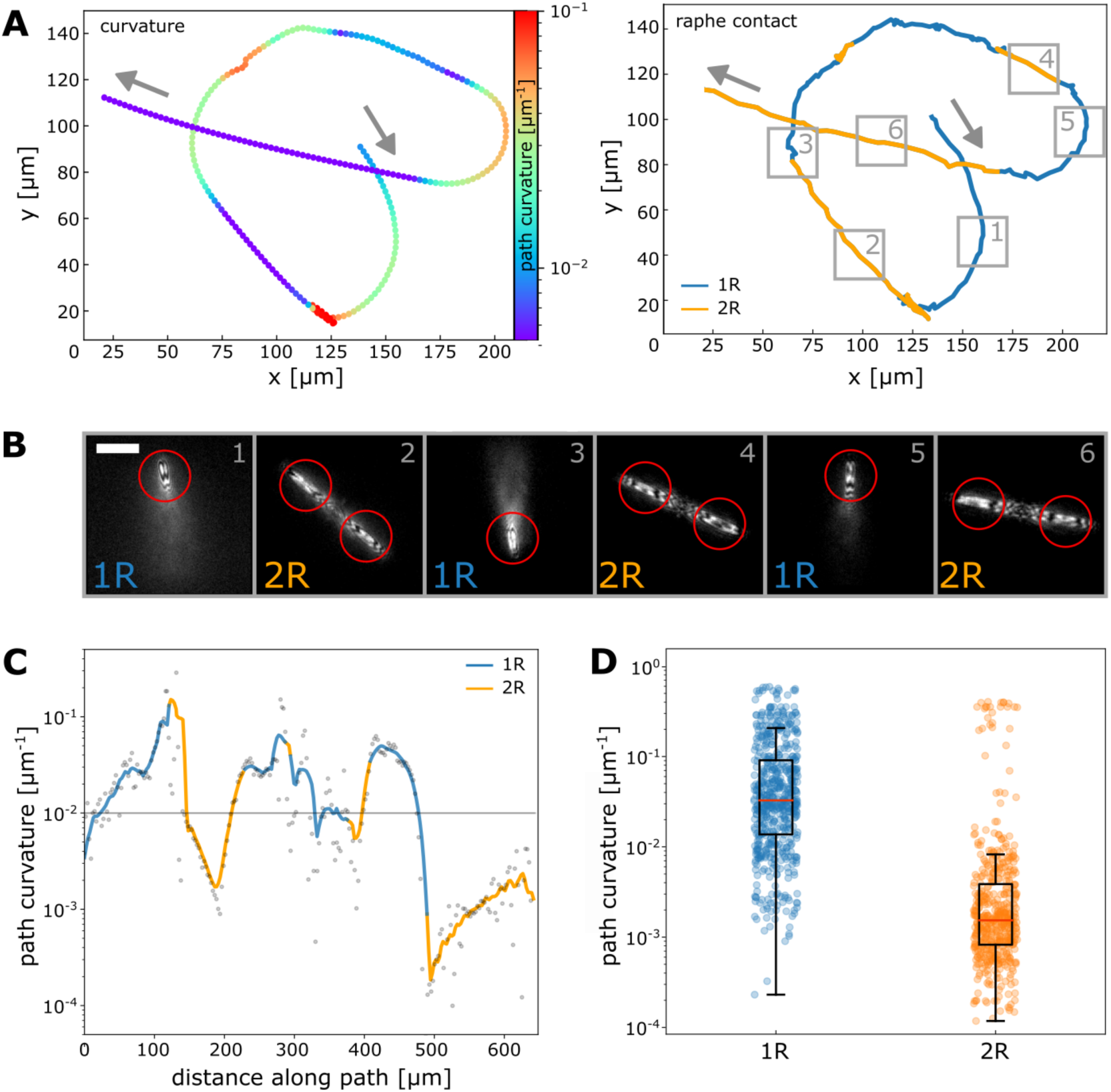
Path curvature in one-raphe and two-raphe contact gliding using interference reflection microscopy (IRM). **A** (left) Trajectory of a single *C. australis* cell showing substantial changes in path curvature (color coded). Grey arrows indicate direction of travel. (right) The same trajectory color-coded by the number of raphes in contact with the substrate during gliding (one-raphe contact gliding in blue, two-raphe contact gliding in orange). **B** IRM images from regions of interest along the trajectory in A, showing the switching between one-raphe and two-raphe contact gliding. Red circles indicate tracked raphes used to discriminate between one- and two-raphe contact gliding. The tracking data of the trailing raphe was used to generate the trajectories in A and B as it was in the present example in constant contact with the substrate. Scale bar is 20 µm. **C** Path curvature over distance along path of the single-cell trajectory in A, with segments of one-raphe contact gliding (blue) and two-raphe contact gliding (orange). **D** Quantification of path curvatures from six different single-cell trajectories displaying substantial changes in path curvature (average cell size of 40µm), pooled by one-raphe (1R) and two-raphe (2R) contact gliding.

### Path curvature is cell size-dependent according to an altered raphe morphology

To validate if changes in raphe morphology indeed lead to predictable alterations in the diatoms’ path curvature, we made use of the fact that *C. australis* undergoes asexual reproduction, resulting in progressively smaller cell sizes with each round of cell division (Christensen, 1991). Analyzing the raphes in SEM images of 15 cell walls from monoclonal cell cultures with bimodal size distribution (average cell size of 36.1 µm and 73.5 µm) revealed that, while the general shape of the raphes remained conserved, the raphes of smaller cells exhibited more bending per unit length than the raphes of the larger cells (**Figure 5A**). Using the procedures described above to predict path curvature from raphe curvature, we accordingly found, that halving the cell size increased the median predicted path curvatures in both one-raphe and two-raphe contact gliding modes by factors of about three (from 0.05 to 0.15 for 1R and 0.0014 to 0.0035 for 2R1 and 2R2, **Figure 5B)**. This predicted cell size-dependent path curvature was confirmed in a population of cells with a similar bimodal size distribution, where a two-fold reduction in cell size led to an about five-fold increase in average path curvature (**Figure 5C and D**). Importantly, our IRM data shows, that the transitions between one-raphe and two-raphe contact gliding, along with its impact on path curvature, remained consistent regardless of cell size (**Supplementary Figure S5**). Taken together, these findings demonstrate that the path curvature of *C. australis* trajectories can be explained by the local curvature of the raphes and the dynamic switching between one-raphe and two-raphe contact gliding, even across a wide range of cell sizes.

**Figure 5:**
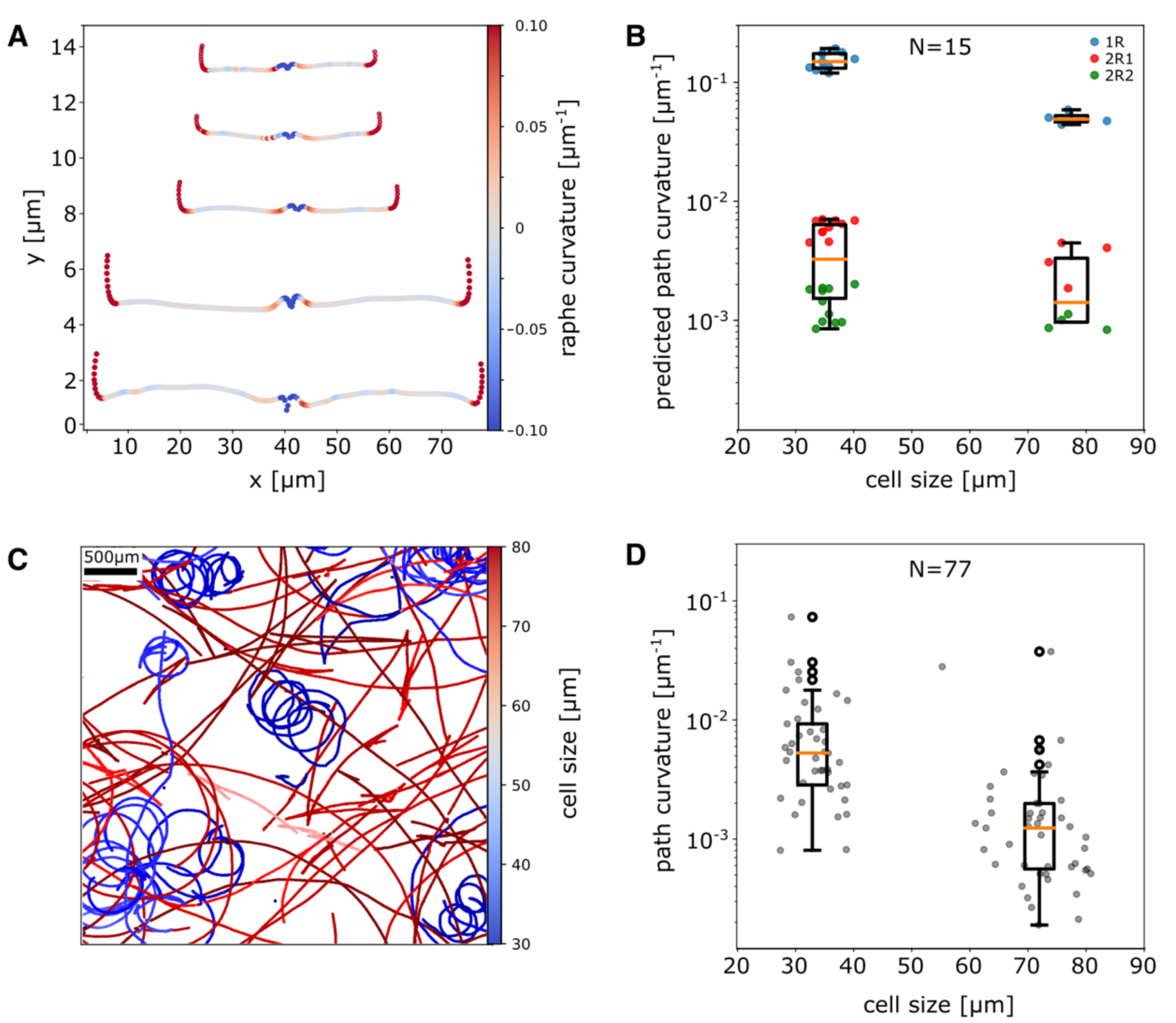
Dependence of raphe and path curvature on cell size. **A** Examples of 5 segmented raphes from SEM images of a *C.australis* cell population with a bimodal size distribution. Raphes are color-coded for curvature with the top three from cells of size ∼36 µm and the bottom two from cells of size ∼78 µm. **B** Predicted average path curvatures for the different modes of gliding, based on the analysis of local raphe curvatures from the same cell population with bimodal cell-size distribution as in **A** (average cell sizes 36.1 µm and 76.5 µm, N = 15 cells). **C** Trajectories of *C. australis* cells from a population with a similar bimodal size distribution as in A. Cells were tracked over one hour (acquired at 1 fps) in a single field of view (3.2 mm x 3.2 mm). Trajectories are color coded by the size of the respective cell. **D** Experimentally observed average path curvatures for each path in C, sorted by cell size (average cell sizes of 32.5 µm and 73.2 µm, N = 77 cells).

## Discussion

In this study we have explored the origin of the dynamic path curvature observed in the trajectories of the motile diatom *C. australis*. While numerous studies have quantified diatom motility characteristics through single-cell tracking and statistical analysis in the past (Edgar, 1979; Edgar & Pickett-Heaps, 1984; Gutiérrez-Medina et al., 2014; Murase et al., 2011; Murguía et al., 2015; Bondoc-Naumovitz et al., 2024), a mechanistic understanding of how diatoms dynamically modify their path shape has remained largely unexplored. Our results show, that the ability of cells to tune path curvature is two-fold. First, small to moderate continuous changes in path curvature may be achieved by shifting the sections where traction for motility is produced along the length of the raphe. This is in line with Harbich’s (2021) hypothesis that the local curvature of the raphe segment in contact with the substrate governs the path shape of a gliding diatom (Harbich, 2021). Secondly, we found that large and abrupt changes in path curvature are achieved by switching between one-raphe and two-raphe contact gliding. It is this dual mechanism, which we term Dynamic Raphe Switching (DRS) that enables directional flexibility across a wide range of temporal and spatial scales, linking geometry of the raphe system to the observed cell size-dependent path curvatures. In contrast to other well-studied motility systems that rely on cell deformation or external appendages, DRS is unique, as it allows for directional flexibility of rigid cells lacking external appendages.

Despite the conserved geometry of the *C. australis* raphe system, the trajectories of similarly sized cells, or even the same cell observed at different times, can vary dramatically. Towards understanding this phenomenon, DRS quantitatively predicts path curvature from local raphe geometries and provides a qualitative and quantitative explanation for the differences in trajectory shapes observed in our experiments. For instance, large trajectory radii - much greater than the cell length - originate from two-raphe contact gliding where both raphes contribute to force generation (2R2), reflecting the combined influence of local raphe curvature in both substrate-contacting segments and their spatial separation. Conversely, abrupt, tight turns with radii comparable to the cell length occur when cells switch from two-raphe to one-raphe contact gliding. Sigmoid paths, another intriguing phenomenon, likely result from cells shifting activity along the raphe system between sections with predominantly positive curvature to sections with predominantly negative curvature. This would result in alternating anti-clockwise and clockwise turns in the trajectory, leading to a sigmoid-shape path segments that we occasionally observed (**Supplementary Movie S9)**. Interestingly, upon directional reversals of cell motility, the orientation of curvature mostly stays the same, suggesting, that either (i) the force generating machinery (intracellular myosins) switches direction along a fixed raphe segment (similar to a car going back and forward with a fixed steering wheel) or (ii) the active zone switches from one raphe to the other. In the latter case, the bilateral symmetry of the raphe system conserves the orientation and magnitude of path curvature. Both behaviors have been observed in the literature (Harbich, 2021; Zhang et al., 2024) and in our IRM experiments (**Supplementary Movie S8**). These mechanisms underpin the stereotypical zig-zag and circular trajectories that are characterized by long runs of constant curvature commonly observed in diatoms (Edgar & Pickett-Heaps, 1984a; Gutiérrez-Medina et al., 2014; Murase et al., 2011). Additionally, our model shows, that the spectrum of path shapes originates from the local raphe geometry, which allows for path curvature to be preserved over distances much greater than the cell size and even upon directional reversals. This is in contrast to other theories, suggesting that propulsion is achieved through hydrodynamic mechanisms, that fail to explain how path shape can be preserved over long distances (Drum & Hopkins, 1966; Singh et al., 2023).

The distinct modes of gliding in our DRS model imply a multimodal distribution of path curvatures on a single-cell level, for which we find strong evidence in our single-cell tracking data (**Supplementary Figure S6**). Furthermore, the transition between two- and one-raphe contact gliding may also account for the observed reduction in velocity with increasing path curvature, as one-raphe contact gliding relies on force generation from only half of the raphe system. These insights provide a deepened understanding of how raphe geometry and force distribution govern the diverse motility patterns of diatom cells.

The proposed DRS motility mechanism relies on the assumption that force production along the raphe system can be heterogeneously distributed. However, the mechanisms by which diatoms regulate the spatial location of force production remains unclear. Recent studies demonstrated that three raphid diatom-specific myosins exhibit coordinated movement during ‘smooth-sustained’ gliding, as well as counteracting movements during pausing or very slow cell movement (Davutoglu et al., 2024). Despite these findings, correlating the movements of these myosins over extended trajectories, such as those described in this study, has yet not been achieved and is subject for future studies. Considering that diatom motility relies on a complex interplay of intracellular force generation by myosin motors, the secretion of adhesive mucilage strands through the raphe, and adhesion to a substrate, it is conceivable that cells exert spatial control over the distribution of raphe-associated myosin motors or regulate adhesive secretion to tune their directional movement. By shifting the location of force generation or adhesive secretion, cells may thus continuously adjust their direction of travel based on the underlying local raphe geometry. Alternatively, selectively halting mucilage secretion in one raphe could facilitate its detachment from the substrate. For rapid and pronounced changes in path curvature, we observed that *C. australis* cells lift and detach one half of their raphe system from the substrate. This phenomenon has been described also for other diatom species by side-view imaging of gliding cells, especially upon directional reversals (Apoya-Horton et al., 2006; Harbich, 2021; Zhang et al., 2024). This lift-up process requires significantly reduced traction in one raphe, allowing the other to dominate motility. Such detachment could result from pronounced asymmetries in intracellular myosin distribution, creating force imbalances that sever adhesive strands in one half of the cell. Force measurements of motile diatoms showed that single cells can produce forces of up to hundreds of pico-Newton (Zheng et al., 2023), equivalent to the coordinated action of hundreds of myosins and enough to sever secreted mucilage strands (Gutiérrez-Medina et al., 2022). However, the precise mechanism linking EPS secretion and diatom cell motility remains an open question.

In our study, we omitted the role of the adhesive mucilage strands to simplify the system, thereby allowing us to quantitatively correlate raphe contact zones with path curvature. Nevertheless, the DRS model is consistent with the AMC model for diatom gliding, as the directional control predicted by the DRS model relies on a mechanical coupling between the intracellular force-generating machinery and the extracellular adhesive strands. That said, the actual site of force production might deviate from where the raphe contacts the substrate due to the length and elastic properties of the adhesive mucilage strands (Higgins et al., 2003; Wang et al., 2000; Zackova Suchanova et al., 2023). Furthermore, the distribution and mechanical properties of the adhesive mucilage, which contribute to the jerky movement of diatoms on short timescales (Gutiérrez-Medina et al., 2022), likely play an important role in shaping motility patterns and even mediate cooperative motility (Zheng et al., 2023). To further investigate the relationship between force generation and substrate traction, techniques such as traction force microscopy combined with fluorescent mucilage staining could provide valuable insights into how mucilage secretion and adhesion influence transitions between one- and two-raphe contact gliding.

The ability of diatoms to dynamically adjust their path shape is crucial for their survival as it allows the cell to follow gradients of light and nutrients and avoid desiccation (Cooksey & Cooksey, 1988; Duchêne et al., 2023; Nultsch, 1971; V. Bondoc et al., 2019). In particular their photoprotective response to varying wavelengths and light intensities allows diatoms to actively explore their light environment and avoid damaging high-intensity light through directed motility (S. A. Cohn, 2001; Morelle et al., 2024). While we have demonstrated that *C. australis* is able to continuously adjust their direction of travel by DRS under isotropic conditions, it remains to be investigated how diatoms adapt their motility in response to anisotropic conditions and environmental gradients.

Quantitative analysis of motility patterns in different diatom species suggest, that many diatoms move in circular random-walk fashion, optimized for biofilm formation and nutrient foraging (Gutiérrez-Medina et al., 2014; Hu et al., 2020). Circular or helical paths have been observed during chemotaxis of other fast moving cells, such as sperm, allowing for an effective sensing length-scale much larger than the cells’ size (Alvarez et al., 2014). Through periodic modulation of their path curvature, sperm cells can measure the chemo-attractant concentration at different spatial positions and robustly follow chemical gradients in complex habitats (Friedrich & Jülicher, 2007). However, whether diatoms employ a similar strategy to optimize their motility in response to environmental gradients will be a focus of future research. Moreover, the molecular and cellular mechanisms underpinning the precise spatial and temporal sensing of environmental cues required for effective adaptation, remain unclear.

Recent work by Helliwell et al. (2019) identified a novel class of diatom-specific ion channels, termed EukCats, which are permeable to both Na^+^ and Ca^2+^ ions (Helliwell et al., 2019) and play a key role in environmental sensing, cellular signaling and Ca^2+^ dependent gliding motility. Additionally, transient increases in cytosolic Ca^2+^ concentrations near the cell apex during directional reversals in response to light further underscore the importance of Ca^2+^ in the intracellular signaling in diatom gliding (McLachlan et al., 2012). Future research could investigate how intracellular myosin dynamics and transitions between one-raphe and two-raphe contact gliding are regulated under anisotropic conditions, allowing diatoms to execute effective taxis and optimize their movement in response to environmental gradients.

Motile diatom species typically inhabit shallow waters and intertidal zones, where motility is critical to avoid desiccation and/or being buried under sediments that are constantly deposited by waves and currents. As such, their natural substrates (i.e. mud, sand, rocks, algae) are highly complex, three-dimensional (3D) and not straight forward to reconstitute in a laboratory setting. Consequently, studying diatom motility on the homogenous, two-dimensional (2D) surface of a coverslip might hence not fully capture the diversity of motility patterns found in their natural habitats. Despite these limitations, analyzing diatom motility on 2D substrates provides a simplified system, enabling us to uncover general mechanisms. Recent studies have attempted to mimic complex 3D natural environments by investigating the motility of diatom populations moving among sand grains, ice and cryolite crystals (Harper, 1969; Consalvey et al., 2004; Zhang et al., 2024; Bondoc-Naumovitz et al., 2024). These studies revealed that controlled 2D assays successfully capture key features of diatom movements and serve as a reliable predictor of their dispersal mechanisms. Over the past decade, advances in techniques for studying cell migration in 3D environments have provided new insights into processes such as parasite infection and cancer progression (Stadler et al., 2022; Brüning-Richardson & Kirby, 2024). Applying similar approaches to diatoms could reveal novel aspects of their motility. Transitioning from 2D to 3D would allow diatom cells greater freedom to adjust their path shape, as bead experiments suggest that all four raphes can function simultaneously (Edgar & Pickett-Heaps, 1984). Taken together, our findings address the long-standing question of how diatoms achieve directional flexibility in their movement despite the static geometry of their raphe system. We show that dynamic switching of cell-substrate contact sites allows gliding diatoms to modulate the curvature of their paths. This novel mechanism advances our understanding of diatoms being able to adapt their motility according to ever-changing conditions in their complex and dynamic habitats.

## Materials and Methods

### Cell culturing

*Craspedostauros australis* (Cox, CCMP3328) was grown in artificial seawater medium EASW (Harrison et al. 1980) in T25 cell culture flasks (Greiner, 690195) at 20°C under a cool white lamp (OSRAM LUMILUX cool daylight L36W/865) at a light intensity of 40 and 60 μmol photons·m^-2^ ·s^-1^ (irradiance of ∼3W/m^2^) Cultures were kept under a 14/10h light/dark cycle and sub-cultured every seven days to maintain the culture in a logarithmic growth phase, as described in (Poulsen et al., 2023).

### Microchamber assembly

Microchambers for microscopy of motile cell populations were assembled by attaching a 24×60mm dichlorodimethylsilane-coated (DDS) coverslip onto an Ibidi sticky slide mount (Ibidi, product 80828) (Gell et al., 2010). The DDS coating on the coverslip rendered the glass surfaces hydrophobic and allowed for long uninterrupted runs of the diatoms. Before the introduction of diatoms, the chambers were flushed with artificial seawater medium EASW to remove any debris before adding an appropriate number of cells to the chamber that would result in about 100 cells per field of view (3.2mm x 3.2mm). Higher cell concentrations were avoided as these lead to frequent cell-cell encounters which might affect the shape of their trajectories.

### Brightfield microscopy and cell motility assessment

For the single-cell motility data presented in this study, *C. australis* cells were transferred from the culture flask to a freshly prepared microchamber and imaged at 22 °C using a Nikon Ti-2 microscope equipped with a Nikon Plan Fluor DL 4x 0.13 air objective and a Lumencor SOLA Light engine LED lamp at 2% intensity and additional 50% neutral density filter and a 530/11nm transmission filter in the light path (irradiance at sample of 4.6 W/m^2^ = 20 µmol photons /m^2^s). Time-lapse videos were acquired using a pco.edge 4.2 sCMOS camera with a spatial resolution of 1.625 µm/ pixel and a temporal resolution of one frame per second at 15 ms exposure time over a time-course of one hour using the Nikon NIS-Elements software.

For motility assessment, data was pre-processed in Fiji (https://imagej.net/), converting the movies into binary movies (reducing the dynamic range of the video to two pixel values of 0 for black and 255 for white), using the inbuild threshold detection method in ImageJ to reduce file-size. Tracking was achieved using a threshold detection method in the Fiji plugin Trackmate version 7.13.2, yielding the spatial coordinates for each single cell at every timepoint from which cell trajectories, velocities and path curvature were calculated using a custom-written python script. Traces shorter than 300 s were excluded from the analysis, as we preferred to study the spectrum of single-cell path curvatures on timescales longer than 5 min. The single cell tracking additionally yielded the length of each cell’s long axis (hence cell size), as well as the orientation of each cell’s long axis at every point in time.

### Assessing path curvature, cell velocity and offset angle in diatom trajectories

From the x-y-t datasets obtained from our single cell tracking, cell trajectories could be reconstituted and motility parameters extracted for each cell and point in time, using a custom-written python script.

To quantify the path curvature *c,* the raw x-y datasets were first pre-processed to smoothen the trajectories using a spline interpolation function. Next, the interpolated datapoints were equally spaced, conserving the total number of datapoints. Path curvature *c* was then computed by analyzing the rate of change of the tangent component of the velocity vector as described in (Lady, 2000):

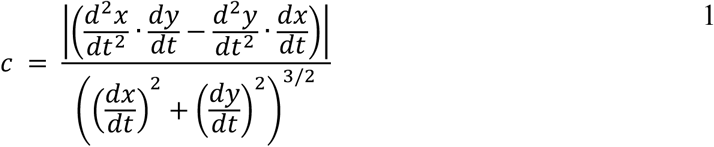

Cell velocity along the path *v* was obtained by first calculating the displacement in the interpolated x and y datasets between two subsequent timepoints and then taking their geometric average divided by the temporal difference between the timepoints:

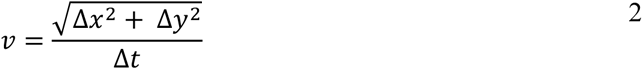

The cell offset angle α was obtained by calculating the absolute difference of the orientation of the cell’s long axis and the direction of travel <π. The orientation of each cell’s long axis for each point in time was obtained from our tracking data as it is a standard return for the threshold detection method in Trackmate (Tinevez et al., 2017). The direction of travel was then computed as follows:

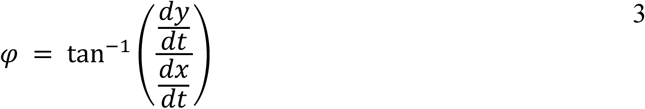

As the time resolution in our long-term single cell tracking data was limited to 1 fps, path curvatures, cell velocities, and offset angles were averaged over 4 µm along the path, using a rolling averaging function. The size of this averaging window was chosen based on the average size of the contact site in our IRM data and maintained for the analysis of raphe curvature, to ensure a similar resolution.

### Scanning electron microscopy (SEM) imaging of diatom cell walls

To isolate diatom cell walls, 200 mL of *C. australis* cell culture (that was used to perform the experiments for data shown in Figure 2) were grown to a density of ∼10^5^ cells mL^−1^. Cells were then harvested using a centrifugation of 5 min at 2,000 xg using a Beckman-Coulter Heraeus Biofuge Stratos centrifuge. The pellet was resuspended in 10 mL extraction buffer (2% SDS, 100 mM EDTA pH 8) and continuously shaken at 55° for 1 hour to solubilize intracellular material. Diatom cell walls were then washed three times by pelleting using a 5 min centrifugation step at 2,000 xg and resuspending in with 2 mL 10 mM EDTA pH 8. Finally, the cleaned cell walls were washed in acetone, followed by centrifugation and resuspension in water. The suspension was then transitioned to 100% ethanol through a series of centrifugation-resuspension steps, with incremental ethanol concentration increases (20%, 40%, 60%, 80%, and 100%). To prevent the collapse of cell walls and preserve their 3D morphology, water-free diatom cell walls were then critical-point dried (using a Leica CPD 300 and mounted onto resin pads for sputter coating with platinum using a Baltec SCD 050 instrument and argon process gas (40 mA, 40 s). SEM images of diatom cell walls were taken using a JSM 7500F field emission scanning electron microscope (Jeol) at an acceleration voltage of 5 kV at the Electron Microscopy core facility at the CMCB Technology Platform at TU Dresden.

### Analysis of the local raphe curvature to predict path curvatures

SEM images of diatom cell walls were preprocessed in Fiji to enhance contrast. The raphes were then manually traced using the segmented line tool, and the extracted x-y coordinate data was used to reconstruct raphe shapes and analyze their curvature with a custom Python script. To ensure uniform spacing of data points, we applied spline interpolation to the x-y datasets of each raphe pair, creating a new dataset of equally spaced points with 10 points per µm (**Supplementary Figure S3A**). Following the model introduced by (Harbich, 2021), we assume that traction is generated only along short sections of the raphe. Based on this, curvature was then analyzed using Equation 1 within 4 µm moving windows that were moved point by point (at a resolution of 10 points per µm) along the raphe. However, unlike path curvature calculations, we allowed for both positive and negative curvatures, as raphes exhibit both clockwise and counterclockwise curvatures, which may cancel each other out and result in path segments with low curvature. To predict path curvature from local raphe curvature, it was necessary to determine which regions of the raphe could establish substrate contact, either individually or simultaneously. Using SEM images of *C. australis* cell walls, we assessed which parts of the raphe could come into proximity with the substrate. According to our model, only raphe regions in direct contact with the substrate contribute to motility. Studying the shape of the C. australis cell wall (**Supplementary Figure S 3B**), we find that these contact areas differ between one-raphe and two-raphe contact gliding.

In the case of two-raphe contact gliding, the cell lies flat on the substrate with both raphes touching the surface. Due to the concave shape of the cell walls at the center, the highly curved central portions of the raphe slit cannot come close to the flat substrate and were therefore excluded from further analysis. Additionally, the highly curved terminal regions of the raphe, located at each end of the cell, are distant from the substrate due to the convex shape at this region of the cell wall. Consequently, only the central portion of each raphe contributes to motility during two-raphe contact gliding, accounting for approximately 50% of the total raphe length in both small and large cells. Since we cannot distinguish, whether forces for motility originated from a single raphe or both raphes simultaneously, we made a further distinction between two-raphe contact with only one active raphe (2R1) and two-raphe contact with both active raphes (2R2). If only one raphe was active, then the resulting path curvature would be dictated by the local raphe curvature along the active stretch. We therefore pooled all curvature values from the central 50% contact zones and obtained a predicted distribution for path curvatures during 2R1 gliding. For the case that both raphes would be active and contributed to path curvature during two-raphe gliding, predicted path curvatures were calculated by fitting a circle through all possible combinations of the 4 µm contact sites on the left and right raphes. The radius of the fitted circle was then used to compute predicted path curvature as c = 1/r, leading to the distribution of predicted curvatures shown in Figure 3D. This indicates that even when both raphes generate traction simultaneously, the cell retains some directional flexibility, allowing for mild variations in path curvature, which were frequently observed in our tracking datasets.

In the case of one-raphe contact gliding, we assumed that one of the two raphes loses traction and lifts up, i.e. away from the substrate. This allows the highly curved terminal regions of the remaining raphe to establish contact with the substrate and contribute to gliding motility and path curvature. Conversely, the central raphe regions cannot make contact during one raphe contact gliding, as this would require the cell to be horizontal to the substrate. Based on geometric considerations from our SEM images (**Supplementary Figure S3B and C**), we estimated that only the terminal 30% of each raphe could contribute to motility during one-raphe contact gliding (**Supplementary Figure S3C**). In these terminal regions, we assumed that traction occurs along 4 µm segments within the contact zones. These 4 µm are based on observations in our later IRM experiments, that revealed typical contact zones of sizes 3 – 10 µm with a median of 4 µm. We thus analyzed the average local raphe curvature within each 4 µm segment for both the left and right raphes independently. We excluded the final 1 µm of each terminal raphe slit, as previous TEM experiments indicated that this region is filled with solid silica and therefore most likely cannot transmit forces (Cox, 1999).

### Interference reflection microscopy (IRM) of gliding diatoms

We use IRM microscopy to determine the number and location of raphe segments where diatoms establish contact with the substrate in concert with the curvature of the cells’ path. To do this we created microchannels by attaching DDS-coated cover slips to Ibidi sticky slides mount (Ibidi, product 80828) and filled the obtained chambers with EASW medium. Prior to imaging, we set up a Nikon Ti-2 microscope for IRM by removing the upper filter and inserted a 50/50 mirror in the lower position filter cube where the dichroic mirror is located, optimizing the incident angle of the light in the Nikon Ti2-LAPP TIRF module for IRM microscopy by closing the aperture diaphragm to approximately 75% (100% is fully open) and installing a Nikon CFI Apochromat 60x 1.49 TIRF oil objective (Mahamdeh & Howard, 2019). As a light source we used the green 561nm line from a LED Lumencor Spectra Light Engine (Lumencor) at an intensity of 1.5% to irradiate the sample from below in the inverted microscope (irradiance at sample of 12 W/m^2^ = 53 µmol photons/m^2^s). The resulting signal was recorded using a pco.edge 4.2 sCMOS camera at 10 ms exposure time with a 2×2 binning of the data to reduce file-size. To increase the limited field of view of a 60x objective, a tiling of 4×4 or 3×3 field of views was used, resulting in a temporal resolution of 4.16 s and 2.56 s per frame respectively. Videos were recorded over 30 min and up to 60 min and later stitched together using the Fiji grid collection stitching plugin with a 15% overlap between the tiles. The position of substrate contacts for each cell were obtained using the Fiji plugin Trackmate v7.13.2. The inbuilt Difference of Gaussian detection mode allowed us to track the position of each contact individually and reconstitute the cells trajectory from the part of the raphe that was in constant contact with the substrate. From the obtained x-y-data, the diatoms trajectory, path curvature and the number of contacts in each frame were extracted using a custom-written python script. The number of contact sides were then used to color-code the cells’ trajectory to and further discriminate path curvatures between one- and two-raphe contact gliding.

## Data, Materials, and Software Availability

All raw and analyzed data, as well as the analysis scripts used to generate the figures in this research article will be made available using Zenodo.

## Acknowledgements

We thank Corina Bräuer and Jennifer Klemm for technical support with cell culture and sample preparation. We thank Dr. Thomas Kurt from the Electron Microscopy Facility of CMCB Technology Platform at TU Dresden for performing critical point drying, sample mounting and SEM imaging of the diatom cell walls. We thank Nils Kröger and Metin Davutoglu as well as all members of the Diez and Kröger labs for helpful discussions and suggestions concerning experiment design and data analysis. We further thank Kirsty Wang for inspiring discussions regarding diatom motility. This research was financed by the Deutsche Forschungsgemeinschaft (DFG, German Research Foundation) under Germany’s Excellence Strategy – EXC 2068 – 390729961 – Cluster of Excellence,Physics of Life of TU Dresden (Nucleation Grant to SD and NP).

## References

Abbriano, R. (2023). Gliding toward new discoveries in diatom adhesion and motility. Journal of Phycology, 59(1), 52–53. 10.1111/jpy.13309

Alvarez, L., Friedrich, B. M., Gompper, G., & Kaupp, U. B. (2014). The computational sperm cell. Trends in Cell Biology, 24(3), 198–207. 10.1016/j.tcb.2013.10.004

Apoya-Horton, M. D., Yin, L., Underwood, G. J. C., & Gretz, M. R. (2006). MOVEMENT MODALITIES AND RESPONSES TO ENVIRONMENTAL CHANGES OF THE MUDFLAT DIATOM *CYLINDROTHECA CLOSTERIUM* (BACILLARIOPHYCEAE) ^1^. Journal of Phycology, 42(2), 379–390. 10.1111/j.1529-8817.2006.00194.x

Bertrand, J. R. (1990). La vitesse de deplacement des Diatomees. Diatom Res., 5, 223–239.

Bertrand, J. R. (2022). Movies about diatoms and their movements. Limnology and Freshwater Biology, 5, 1643–1644. 10.31951/2658-3518-2022-A-5-1643

Bondoc, K. G. V., Heuschele, J., Gillard, J., Vyverman, W., & Pohnert, G. (2016). Selective silicate-directed motility in diatoms. Nature Communications, 7(1), 10540. 10.1038/ncomms10540

Bondoc, K. G. V., Lembke, C., Vyverman, W., & Pohnert, G. (2016). Searching for a Mate: Pheromone-Directed Movement of the Benthic Diatom Seminavis robusta. Microbial Ecology, 72(2), 287–294. 10.1007/s00248-016-0796-7

Bondoc-Naumovitz, K. G., Crosato, E., & Wan, K. Y. (2024). Diversity of motility patterns in benthic diatoms. Biophysics. 10.1101/2024.12.23.630140

Brüning-Richardson, A., & Kirby, C. (2024). Cell Migration in Cancer; Cell Migration in 2D and 3D. In A. Brüning-Richardson & S. Knipp (Eds.), Cell Migration in Development, Health and Disease (pp. 111–137). Springer Nature Switzerland. 10.1007/978-3-031-64532-7_5

Christensen, T. (1991). The diatoms. Biology and morphology of the genera. Taylor & Francis.

Cohn, S. A. (2001). Chapter 13 Photo-stimulated effects on diatom motility. In Comprehensive Series in Photosciences (Vol. 1, pp. 375–401). Elsevier. 10.1016/S1568-461X(01)80017-X

Cohn, S. A., Dunbar, S., Ragland, R., Schulze, J., Suchar, A., Weiss, J., & Wolske, A. (2016). Analysis of light quality and assemblage composition on diatom motility and accumulation rate. Diatom Research, 31(3), 173–184. 10.1080/0269249X.2016.1193058

Cohn, S. A., Halpin, D., Hawley, N., Ismail, A., Kaplan, Z., Kordes, T., Kuhn, J., Macke, W., Marhaver, K., Ness, B., Olszewski, S., Pike, A., Rice, E., Sbarboro, J., Wolske, A., & Zapata, Y. (2015). Comparative analysis of light-stimulated motility responses in three diatom species. Diatom Research, 30(3), 213–225. 10.1080/0269249X.2015.1058295

Cohn, S. A., Spurck, T. P., & Pickett-Heaps, J. D. (1999). HIGH ENERGY IRRADIATION AT THE LEADING TIP OF MOVING DIATOMS CAUSES A RAPID CHANGE OF CELL DIRECTION. Diatom Research, 14(2), 193–206. 10.1080/0269249X.1999.9705466

Cohn, S. A., & Weitzell, R. E. (1996). ECOLOGICAL CONSIDERATIONS OF DIATOM CELL MOTILITY. I. CHARACTERIZATION OF MOTILITY AND ADHESION IN FOUR DIATOM SPECIES ^1^. Journal of Phycology, 32(6), 928–939. 10.1111/j.0022-3646.1996.00928.x

Consalvey, M., Paterson, D. M., & Underwood, G. J. C. (2004). THE UPS AND DOWNS OF LIFE IN A BENTHIC BIOFILM: MIGRATION OF BENTHIC DIATOMS. Diatom Research, 19(2), 181–202. 10.1080/0269249X.2004.9705870

Cooksey, B., & Cooksey, K. E. (1988). Chemical signal-response in diatoms of the genus *Amphora*. Journal of Cell Science, 91(4), 523–529. 10.1242/jcs.91.4.523

Cox, E. (1999). Craspedostauros gen. Nov., a new diatom genus for some unusual marine raphid species previously placed in Stauroneis Ehrenberg and Stauronella Mereschkowsky. European Journal of Phycology, 34(2), 131–147. 10.1080/09670269910001736192

Davutoglu, M. G., Geyer, V. F., Niese, L., Soltwedel, J. R., Zoccoler, M. L., Sabatino, V., Haase, R., Kröger, N., Diez, S., & Poulsen, N. (2024). Gliding motility of the diatom Craspedostauros australis coincides with the intracellular movement of raphid-specific myosins. Communications Biology, 7(1), 1187. 10.1038/s42003-024-06889-w

Drum, R. W., & Hopkins, J. T. (1966). Diatom locomotion: An explanation. Protoplasma, 62(1), 1–33. 10.1007/BF01254629

Duchêne, C., Bouly, J.-P., Karlusich, J. J. P., Sellés, J., & Bowler, C. (2023). Diatom phytochromes integrate the entire visible light spectra for photosensing in marine environments. bioRxiv. 10.1101/2023.01.25.525482

Edgar, L. A. (1979). Diatom locomotion: Computer assisted analysis of cine film. British Phycological Journal, 14(1), 83–101. 10.1080/00071617900650111

Edgar, L. A. (1983). Mucilage secretions of moving diatoms. Protoplasma, 118(1), 44–48. 10.1007/BF01284745

Edgar, L. A., & Pickett-Heaps, J. (1984). Diatom locomotion. Progress in Phycological Research.

Friedrich, B. M., & Jülicher, F. (2007). Chemotaxis of sperm cells. Proceedings of the National Academy of Sciences, 104(33), 13256–13261. 10.1073/pnas.0703530104

Gell, C., Bormuth, V., Brouhard, G. J., Cohen, D. N., Diez, S., Friel, C. T., Helenius, J., Nitzsche, B., Petzold, H., Ribbe, J., Schäffer, E., Stear, J. H., Trushko, A., Varga, V., Widlund, P. O., Zanic, M., & Howard, J. (2010). Microtubule Dynamics Reconstituted In Vitro and Imaged by Single-Molecule Fluorescence Microscopy. In Methods in Cell Biology (Vol. 95, pp. 221–245). Elsevier. 10.1016/S0091-679X(10)95013-9

Gutiérrez-Medina, B., Guerra, A. J., Maldonado, A. I. P., Rubio, Y. C., & Meza, J. V. G. (2014). Circular random motion in diatom gliding under isotropic conditions. Physical Biology, 11(6), 066006. 10.1088/1478-3975/11/6/066006

Gutiérrez-Medina, B., Peña Maldonado, A. I., & García-Meza, J. V. (2022). Mechanical testing of particle streaming and intact extracellular mucilage nanofibers reveal a role of elastic force in diatom motility. Physical Biology, 19(5), 056002. 10.1088/1478-3975/ac7d30

Harbich, T. (2021). Some Observations of Movements of Pennate Diatoms in Cultures and Their Possible Interpretation. In S. Cohn, K. Manoylov, & R. Gordon (Eds.), Diatom Gliding Motility (1st ed., pp. 1–31). Wiley. 10.1002/9781119526483.ch1

Harper, M. A. (1969). Movement and migration of diatoms on sand grains. British Phycological Journal, 4(1), 97–103. 10.1080/00071616900650081

Harper, M. A., & Harper, J. F. (1967). Measurements of diatom adhesion and their relationship with movement. British Phycological Bulletin, 3(2), 195–207. 10.1080/00071616700650051

Helliwell, K. E., Chrachri, A., Koester, J. A., Wharam, S., Verret, F., Taylor, A. R., Wheeler, G. L., & Brownlee, C. (2019). Alternative Mechanisms for Fast Na+/Ca2+ Signaling in Eukaryotes via a Novel Class of Single-Domain Voltage-Gated Channels. Current Biology, 29(9), 1503–1511.e6. 10.1016/j.cub.2019.03.041

Higgins, M. J., Molino, P., Mulvaney, P., & Wetherbee, R. (2003). THE STRUCTURE AND NANOMECHANICAL PROPERTIES OF THE ADHESIVE MUCILAGE THAT MEDIATES DIATOM-SUBSTRATUM ADHESION AND MOTILITY^1^. Journal of Phycology, 39(6), 1181–1193. 10.1111/j.0022-3646.2003.03-027.x

Hu, W. S., Huang, M., Zhang, H. P., Zhang, F., Vyverman, W., & Liu, Q. X. (2020). Movement behavioral plasticity of benthic diatoms driven by optimal foraging. bioRxiv. 10.22541/au.157964086.65973830

Jesus, B., Jauffrais, T., Trampe, E., Méléder, V., Ribeiro, L., Bernhard, J. M., Geslin, E., & Kühl, M. (2023). Microscale imaging sheds light on species-specific strategies for photo-regulation and photo-acclimation of microphytobenthic diatoms. Environmental Microbiology, 25(12), 3087–3103. 10.1111/1462-2920.16499

Lady, L. (2000). Curvature. University of Hawaii. https://math.hawaii.edu/∼lee/calculus/curvature.pdf

Lind, J. L., Heimann, K., Miller, E. A., Van Vliet, C., Hoogenraad, N. J., & Wetherbee, R. (1997). Substratum adhesion and gliding in a diatom are mediated by extracellular proteoglycans. Planta, 203(2), 213–221. 10.1007/s004250050184

Mahamdeh, M., & Howard, J. (2019). Implementation of interference reflection microscopy for label-free, high-speed imaging of microtubules.

McBride, M. J. (2001). Bacterial Gliding Motility: Multiple Mechanisms for Cell Movement over Surfaces. Annual Review of Microbiology, 55(1), 49–75. 10.1146/annurev.micro.55.1.49

McLachlan, D. H., Brownlee, C., Taylor, A. R., Geider, R. J., & Underwood, G. J. C. (2009). LIGHT-INDUCED MOTILE RESPONSES OF THE ESTUARINE BENTHIC DIATOMS *NAVICULA PERMINUTA* AND *CYLINDROTHECA CLOSTERIUM* (BACILLARIOPHYCEAE) ^1^. Journal of Phycology, 45(3), 592–599. 10.1111/j.1529-8817.2009.00681.x

McLachlan, D. H., Underwood, G. J. C., Taylor, A. R., & Brownlee, C. (2012). CALCIUM RELEASE FROM INTRACELLULAR STORES IS NECESSARY FOR THE PHOTOPHOBIC RESPONSE IN THE BENTHIC DIATOM *NAVICULA PERMINUTA* (BACILLARIOPHYCEAE) ^1^. Journal of Phycology, 48(3), 675–681. 10.1111/j.1529-8817.2012.01158.x

Milo, R., & Phillips, R. (2015). Cell biology by the numbers. Garland Science.

Morelle, J., Bastos, A., Frankenbach, S., Frommlet, J. C., Campbell, D. A., Lavaud, J., & Serôdio, J. (2024). The Photoprotective Behavior of a Motile Benthic Diatom as Elucidated from the Interplay Between Cell Motility and Physiological Responses to a Light Microgradient Using a Novel Experimental Setup. Microbial Ecology, 87(1), 40. 10.1007/s00248-024-02354-7

Murase, A., Kubota, Y., Hirayama, S., Kumashiro, Y., Okano, T., Mayama, S., & Umemura, K. (2011). Two-dimensional trajectory analysis of the diatom Navicula sp. Using a micro chamber. Journal of Microbiological Methods, 87(3), 316–319. 10.1016/j.mimet.2011.09.006

Murguía, J. S., Rosu, H. C., Jimenez, A., Gutiérrez-Medina, B., & García-Meza, J. V. (2015). The Hurst exponents of Nitzschia sp. Diatom trajectories observed by light microscopy. Physica A: Statistical Mechanics and Its Applications, 417, 176–184. 10.1016/j.physa.2014.09.046

Nultsch, W. (1956). Studien über die Phototaxis der Diatomeen. Arch. Protistenk, 101(1).

Nultsch, W. (1971). PHOTOTACTIC AND PHOTOKINETIC ACTION SPECTRA OF THE DIATOM *NITZSCHIA COMMUNIS*. Photochemistry and Photobiology, 14(6), 705– 712. 10.1111/j.1751-1097.1971.tb06209.x

Poulsen, N., Davutoglu, M. G., & Zackova Suchanova, J. (2022). Diatom Adhesion and Motility. In A. Falciatore & T. Mock (Eds.), The Molecular Life of Diatoms (pp. 367– 393). Springer International Publishing. 10.1007/978-3-030-92499-7_14

Poulsen, N., Hennig, H., Geyer, V. F., Diez, S., Wetherbee, R., Fitz-Gibbon, S., Pellegrini, M., & Kröger, N. (2023). On the role of cell surface associated, mucin-like glycoproteins in the pennate diatom *Craspedostauros australis* (Bacillariophyceae). Journal of Phycology, 59(1), 54–69. 10.1111/jpy.13287

Round, F. E. (1990). The Diatoms: Biology and Morphology of the Genera (Vol. 747). Cambridge University Press.

Singh, S. R. V., Katyal, K., & Gordon, R. (2023). RAPHE: Simulation of the Dynamics of Diatom Motility at the Molecular Level–The Domino Effect Hydration Model with Concerted Diffusion. The Mathematical Biology of Diatoms, 291–342.

Stadler, R. V., Nelson, S. R., Warshaw, D. M., & Ward, G. E. (2022). A circular zone of attachment to the extracellular matrix provides directionality to the motility of *Toxoplasma gondii* in 3D. eLife, 11, e85171. 10.7554/eLife.85171

Tinevez, J.-Y., Perry, N., Schindelin, J., Hoopes, G. M., Reynolds, G. D., Laplantine, E., Bednarek, S. Y., Shorte, S. L., & Eliceiri, K. W. (2017). TrackMate: An open and extensible platform for single-particle tracking. Methods, 115, 80–90. 10.1016/j.ymeth.2016.09.016

V. Bondoc, K. G., Lembke, C., Vyverman, W., & Pohnert, G. (2019). Selective chemoattraction of the benthic diatom *Seminavis robusta* to phosphate but not to inorganic nitrogen sources contributes to biofilm structuring. MicrobiologyOpen, 8(4), e00694. 10.1002/mbo3.694

Wang, Y., Chen, Y., Lavin, C., & Gretz, M. R. (2000). Extracellular matrix assembly in diatoms (Bacillariophyceae). Iv. Ultrastructure of Achnanthes longipes and Cymbella cistula as revealed by high-pressure freezing/freeze substituton and cryo-field emission scanning electron microscopy. Journal of Phycology, 36(2), 367–378. 10.1046/j.1529-8817.2000.99102.x

Zackova Suchanova, J., Bilcke, G., Romanowska, B., Fatlawi, A., Pippel, M., Skeffington, A., Schroeder, M., Vyverman, W., Vandepoele, K., Kröger, N., & Poulsen, N. (2023). Diatom adhesive trail proteins acquired by horizontal gene transfer from bacteria serve as primers for marine biofilm formation. New Phytologist, 240(2), 770–783. 10.1111/nph.19145

Zhang, Q., Leng, H. T., Li, H., Arrigo, K. R., & Prakash, M. (2024). Ice gliding diatoms establish record-low temperature limit for motility in a eukaryotic cell. Biophysics. 10.1101/2024.11.18.624199

Zheng, P., Kumadaki, K., Quek, C., Lim, Z. H., Ashenafi, Y., Yip, Z. T., Newby, J., Alverson, A. J., Jie, Y., & Jedd, G. (2023). Cooperative motility, force generation and mechanosensing in a foraging non-photosynthetic diatom. Open Biology. 10.1098/rsob.230148

