## Supplementary Appendix for "Dynamic switching of cell-substrate contact sites allows gliding diatoms to modulate the curvature of their paths"

**This PDF file includes:**

Figures S1 to S6
Legends for Movies S1 to S9

### Supplementary Figures

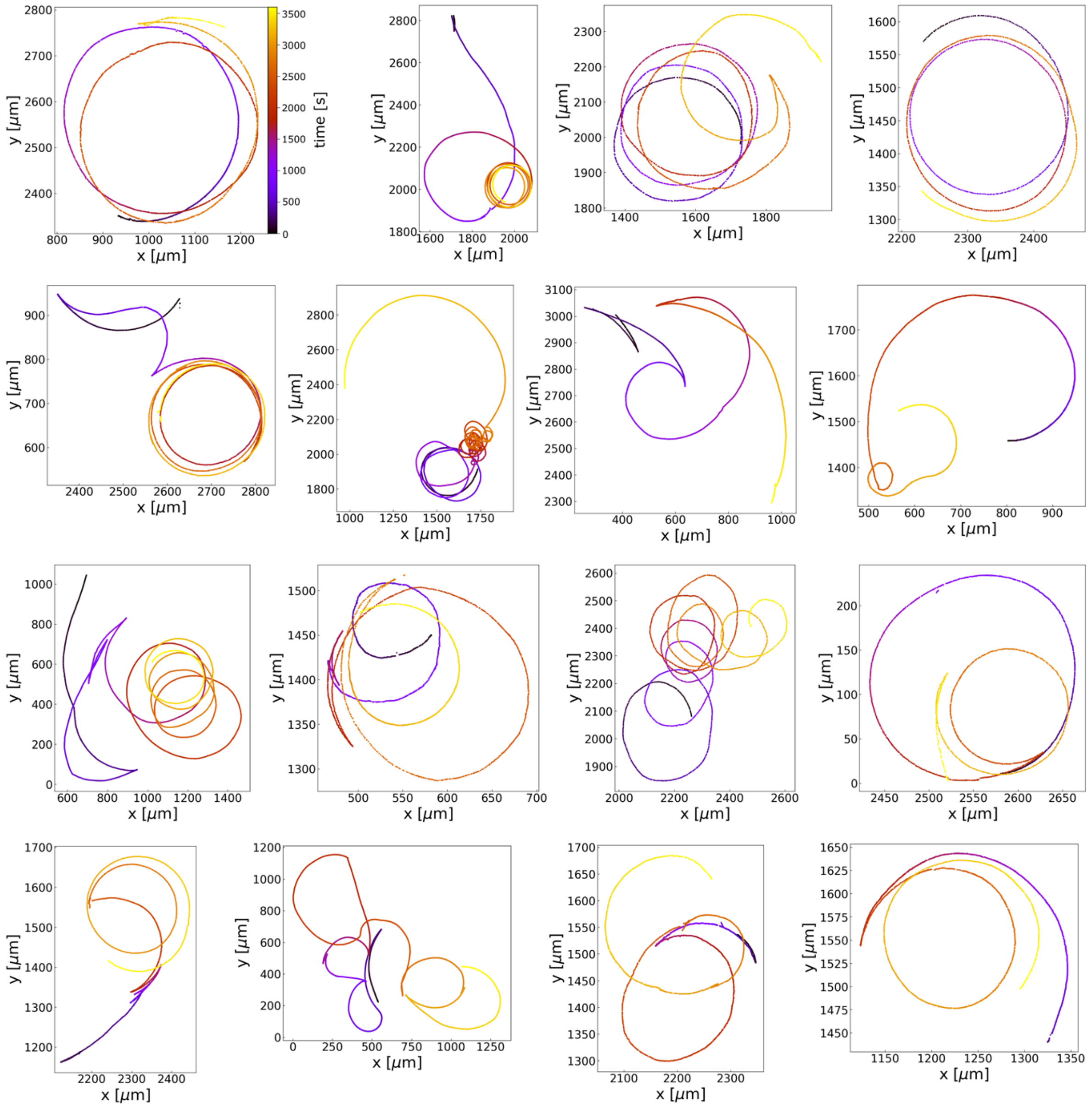


**Supplementary Figure S1: Examples of single-cell trajectories of motile *C. australis*.** Trajectories of individual *C. australis* cells from the population depicted in Figure 1C, tracked at 1 fps over a period of one hour (color-code is time). Cell culture with a homogeneous cell size distribution (average cell size of 25.8 μm +/- 2.0 μm).

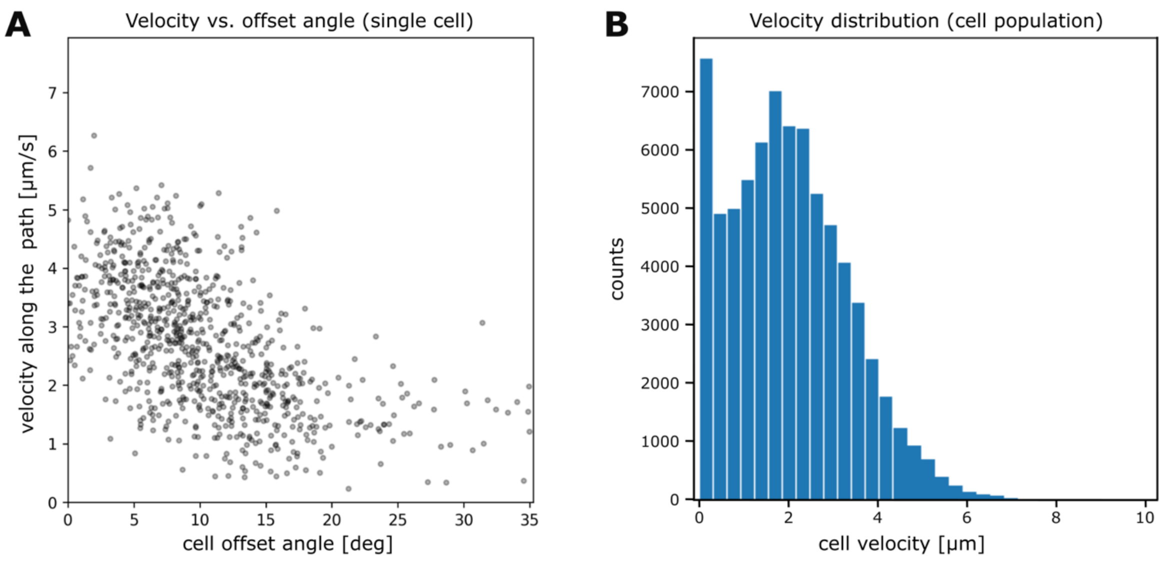


**Supplementary Figure S2: Further quantification of motility parameters. A** Correlation of cell velocity along the path over cell offset angle for the single cell trajectory in Figure 2 A-C. **B** Histogram of cell velocities along the path for the cell population in Figure 2 D-E.

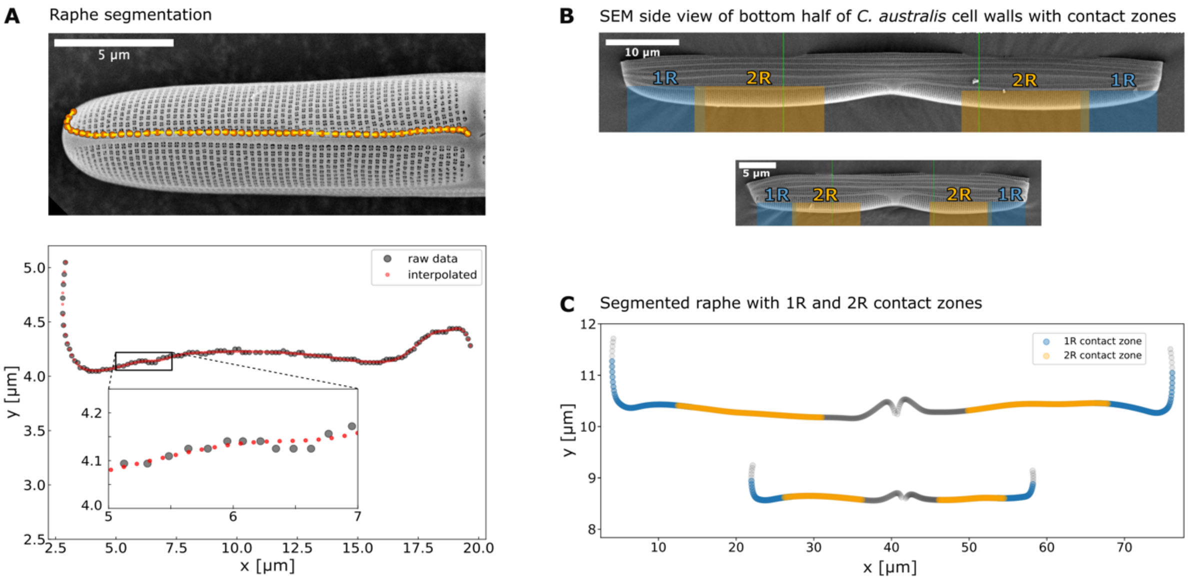


**Supplementary Figure S3: Raphe segmentation and contact zone allocation. A** Details on raphe segmentation and raphe curvature analysis. (top) SEM image of one half of a diatom cell wall with the manually segmented raphe slit (yellow line). (bottom) Plot of the x-y-data obtained from the segmented raphe slit with raw data in grey and interpolated data points in red. To ensure a constant spatial resolution and eliminate fluctuating distances between data points due to manual raphe segmentation, a new set of interpolated points (red dots) is set at an equal spacing with a resolution of 10 points per micrometer. **B** SEM images of the bottom halves of two *C. australis* silica cell walls from two cells of sizes ~76 µm (top) and ~38 µm (bottom) with marked contact zones for one-raphe (blue rectangle) and two-raphe (orange rectangle) contact gliding. **C** Segmented raphes of two cells of similar sizes as in **B**, with raw data in grey and zones for one-raphe and two-raphe contact gliding colored in blue and orange respectively. Note that the terminal 1 µm in each raphe is excluded from analysis, as the terminal raphe fissure is closed and hence most likely does not allow for force transduction.

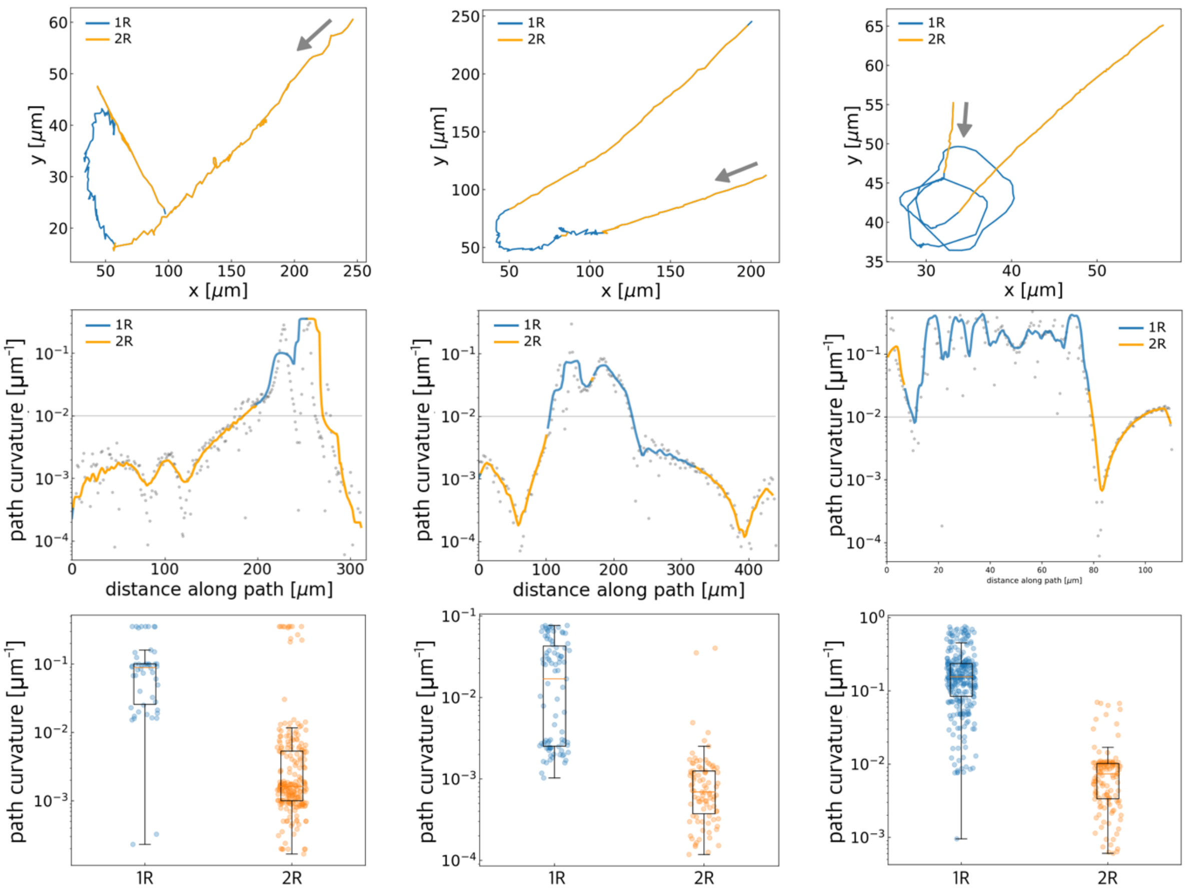


**Supplementary Figure S4: Path curvature analysis during one-raphe and two-raphe contact gliding using interference reflection microscopy (IRM).** (top) Parts of three trajectories of single *C. australis* cells of (average cell size of 40 µm), showing abrupt changes in path curvature, color-coded for the number of raphes in contact with the substrate (detected using IRM) during gliding: one-raphe contact gliding in blue, two-raphe contact gliding in orange. Grey arrows indicate direction of travel. (middle) Path curvature over distance along path of the respective single-cell trajectories with raw data as grey dots and rolling average (over window of 4 µm), color-coded for one-raphe contact gliding (blue) and two-raphe contact gliding (orange). (bottom) Statistical quantification of path curvatures for each trajectory, pooled by one-raphe and two-raphe contact gliding (blue and orange dots respectively).

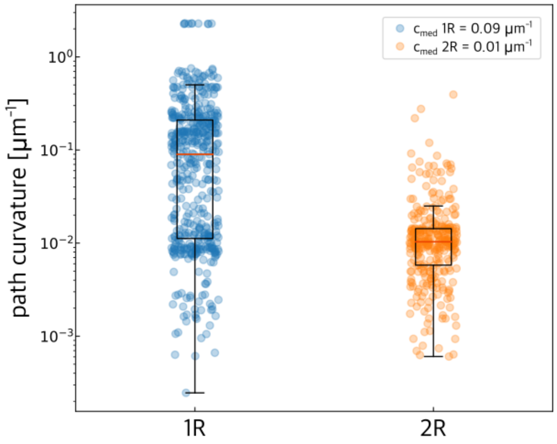


**Supplementary Figure S5: Control for cell size – dependency of dynamic raphe switching (DRS).** Quantification of path curvatures from five different single-cell trajectories showing large, abrupt changes in path curvature from cells only half the size (average cell size of 19 µm) of the cells in Figure 4 and Supplementary Figure S4 (average cell size of 40 µm). Curvature data pooled by one-raphe (1R) and two-raphe (2R) contact gliding (detected by IRM) shows, that the DRS mechanism is independent of cell size and its effect on the ensuing path curvature is conserved.

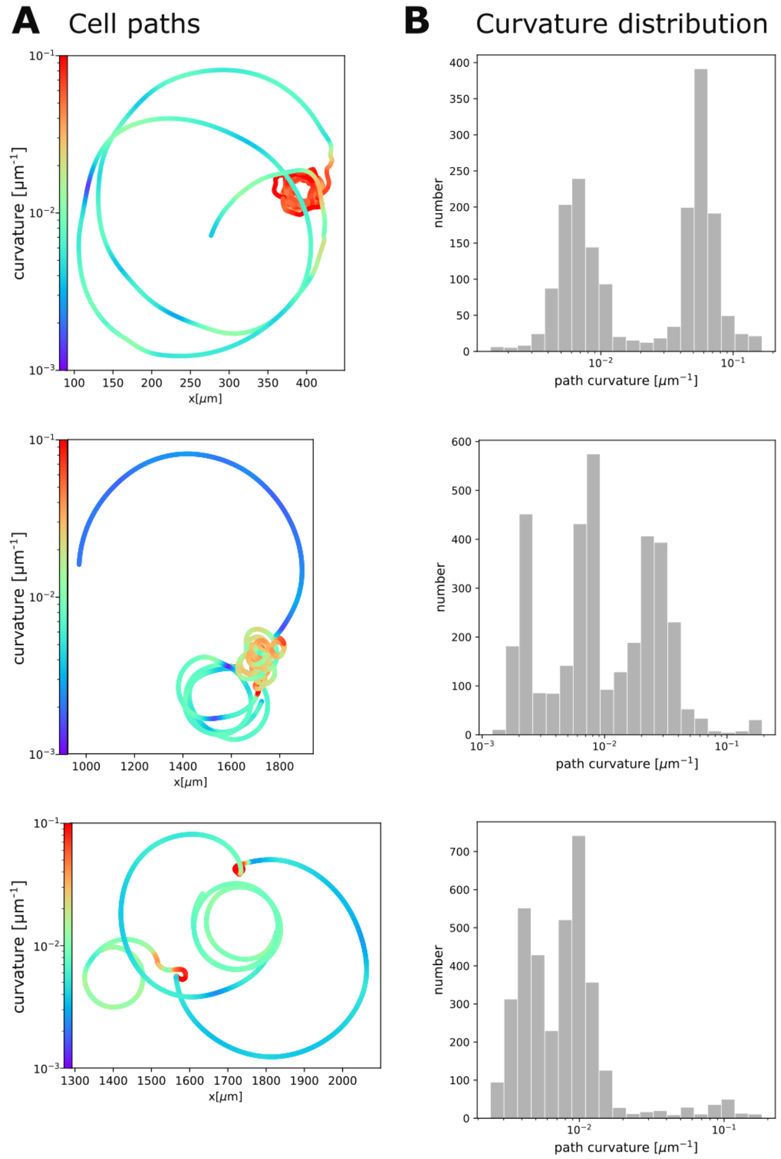


**Supplementary Figure S6: Examples of multi-modal path curvature distributions indicating multiple modes of gliding motility. A** Example trajectories of single *C. australis* cells of sizes 20 µm (top), 25 µm (mid) and 24 µm (bottom), tracked over one hour and color-coded for path curvature. **B** Path curvature distributions for the respective trajectories in A, revealing distinct modes of path curvatures, indicative of distinct modes of gliding motility.

**Legends for movies S1 to S9**

**Supplementary Movie S1:** Example of a tracked cell population of about 80 individual cells in one field of view, imaged at 4x magnification and 1 fps for one hour and tracked using the Fiji-Plugin Trackmate (threshold detection method). Each cell trajectory is displayed in a different color to improve contrast between individual tracks. The raw microscopy data was preprocessed by inverting contrast and creating a binary image to reduce file size.

**Supplementary Movie S2:** Example of an individual diatom trajectory displayed in Figure 1C, imaged at 4x magnification and 1 fps for one hour and tracked with Fiji-Plugin Trackmate (threshold detection method). The raw microscopy data was preprocessed by inverting contrast and creating a binary image to reduce file size.

**Supplementary Movie S3:** Part of an individual motile diatom trajectory used to quantify motility parameters in Figure 2A-E. The cell was imaged at 4x magnification and 1 fps and tracked with Fiji-Plugin Trackmate (threshold detection method). The raw microscopy data was preprocessed by inverting contrast and creating a binary image to reduce file size.

**Supplementary Movie S4:** The same cell as in supplementary Movie S3, but now with the front and back of the cell tracked individually to display the offset between the two trajectories along stretches of high curvature. The raw microscopy data was preprocessed by inverting contrast and creating a binary image to reduce file size. Tracking was done using the Difference of Gaussian detector in the Fiji plugin Trackmate.

**Supplementary Movie S5:** Part of another individual motile diatom trajectory with the front and back of the cell tracked individually to display the offset between the two trajectories along stretches of high curvature. The raw microscopy data was preprocessed using 2x2 binning, inverting contrast and creating a binary image to reduce file size. Tracking was done using the Difference of Gaussian detector in the Fiji plugin Trackmate.

**Supplementary Movie S6:** IRM microscopy of an individual motile diatom cell used to create panels in Figure 4 A-D. Contact-sites between cell and substrate were tracked using the Difference of Gaussian detector in the Fiji plugin Trackmate and colored differently to enhance contrast between individual tracks. The cell was imaged at 60x magnification and 0.25 fps.

**Supplementary Movie S7:** IRM microscopy of another individual motile diatom showing abrupt changes in path curvature in concert with switches from two to one raphe gliding. Contact-sites between cell and substrate were tracked using the Difference of Gaussian detector in the Fiji plugin Trackmate and colored differently to enhance contrast between individual tracks. The cell was imaged at 60x magnification and 0.25 fps.

**Supplementary Movie S8:** Another example of IRM microscopy of an individual motile *C.australis* cell with cell-substrate contact-site tracked using the Difference of Gaussian detector in the Fiji plugin Trackmate and colored differently to enhance contrast between individual tracks. Note that the both halves of the cell alternatingly detach from the substrate during repositioning of the cell body. The cell was imaged at 60x magnification and 0.25 fps.

**Supplementary Movie S9:** Part of an individual diatom trajectory displaying rare sigmoid path shape by changing from a clockwise to a counter-clockwise curve. Note that upon directional reversal, the direction of curvature does not change (continues as counter-clockwise), which is also rare. Cell was imaged at 4x magnification and 1 fps, then tracked with Fiji-Plugin Trackmate. The raw microscopy data was preprocessed by inverting contrast and creating a binary image to reduce file size.
